# Brain activity links performance in science reasoning with conceptual approach

**DOI:** 10.1101/526574

**Authors:** Jessica E. Bartley, Michael C. Riedel, Taylor Salo, Emily R. Boeving, Katherine L. Bottenhorn, Elsa I. Bravo, Rosalie Odean, Alina Nazareth, Robert W. Laird, Matthew T. Sutherland, Shannon M. Pruden, Eric Brewe, Angela R. Laird

## Abstract

Understanding how students learn is crucial for helping them succeed. We examined brain function in 107 undergraduate students during a task known to be challenging for many students – physics problem solving – to characterize underlying neural mechanisms and determine how these support comprehension and proficiency. Further, we applied module analysis to response distributions, defining groups of students who answered using similar physics conceptions, and probed for brain differences linked with different conceptual approaches. We found integrated executive, attentional, visual motion, and default mode brain systems cooperate to achieve sequential and sustained physics-related cognition. While accuracy alone did not predict brain function, dissociable brain patterns were observed when students solved problems using different physics conceptions, and increased success was linked to conceptual coherence. Our analyses demonstrate that episodic associations and control processes operate in tandem to support physics reasoning, offering potential insight to support student learning.

---

New innovations in transforming science education to promote success and broaden participation require an understanding of how students learn. Evidence has shown that learning interventions, both long- and short-term, can be accompanied by lasting, content-related brain changes, suggesting classroom instruction may influence the measurable neural processes by which students consolidate, access, or store information (1, 2). Physics in particular can be a challenging discipline for many students as it requires both a conceptual understanding and recall of physical principles, along with acquisition of procedural skills for solving problems. Neuroimaging studies on physics learning indicate cognition about physical concepts (e.g., velocity, acceleration, force) are encoded into specific neural representations (3), and these representations may change during progressive stages of physics learning (4). Moreover, problem solving is known to engage an extensive frontoparietal central executive network (CEN), both generally across domains of knowledge (5) and specifically regarding physics concepts (6). Collectively, these findings highlight a putatively influential role science learning may have on functional brain architecture and underscore the complexity of neural processes linked with proficiency in physics problem solving.

Insight into the scientific learning process may be gained by considering the obstacles students encounter. A wealth of cognitive science and education research has identified consistent patterns in how students think about physics, with a preponderance of studies focusing on difficulties mastering Newtonian mechanics (7–9). Physics students consistently struggle to learn key concepts and novice students are known to invoke intuitive but incorrect ideas of physical causality when solving problems (10, 11). These misleading conceptions frequently interfere with a student’s ability to successfully acquire new physics knowledge (12) and a broad, but sometimes conflicting, body of literature has attempted to characterize these ideas to support conceptual change across instruction (13–17). One model posits these so-called “folk physics” notions (18, 19) may be implicitly linked to associative memory, with naïve reasoning arising from context-based extrapolations of remembered personal experiences (20). Another describes students’ reasoning as being based on common sense, but weakly organized, physical intuitions (21). Yet another view argues ontological differences in the way students think about physical processes impact how persistent incorrect conceptions are across instruction (14). A contrasting opinion holds that students use ontological categories dynamically and that the range of physics reasoning processes may be better explained by varying levels of coherence (integration of concepts) and robustness (applicability across contexts) in how students build patterns of associations between their existing cognitive resources (e.g., memories, beliefs, facts; (15, 22)). Despite these many models, little is known about the underlying neural processes of how students access, deploy, and attempt to resolve physics conceptions during reasoning. The limited work that has been done on this topic indicates the anterior cingulate cortex (ACC) may be engaged when students view physically causal scenes that conflict with their strongly held intuitions (23). Additionally, episodic, associative, and spatial recall are know to be supported by hippocampal and retrosplenial cortex (RSC) activity (24, 25), and reasoning processes are linked with the dorsolateral prefrontal (dlPFC) and posterior parietal cortex (PPC) activity (5). However, no prior work has identified the specific neural processes that underlie physics reasoning nor any neurobiological differences associated with students different use of incorrect physics conceptions. Such an understanding would inform existing behavioral models and might help us more fully understand how students learn physics.

We acquired functional magnetic resonance imaging (fMRI) data from 107 undergraduate students after the conclusion of a semester of university-level physics instruction. During fMRI, students were presented with questions adapted from the Force Concept Inventory (FCI; (26)), a widely adopted test of conceptual problem solving that presents scenarios of objects at rest or in motion and asks students to choose between a Newtonian solution and several reasonable Non-Newtonian alternatives, each of which mirror common confusions. Physics and baseline perceptual questions (**Fig. S1**) were presented as blocks composed of three sequential view screens (e.g., “phases”): problem initiation in which students viewed text and a figure describing a physical scenario (Phase I), question presentation in which the students viewed a physics question about the scenario (Phase II), and answer selection wherein four possible answer choices were displayed for selection (Phase III). Brain activity across full questions (All Phases), as well as within each phase, was assessed. We then explored putative links between the neural substrates of physics problem solving and accuracy, difficulty, strategy, and student conceptualization of physics ideas. First, we probed for brain-behavior correlations revealed by parametric modulation of the BOLD signal in independent meta-analytically defined *a priori* reasoning and memory-linked regions of interest (ROIs; **Fig. S2**) located in the left dlPFC, ACC, left PPC, left hippocampus, and RSC, and across the whole brain. Second, because student response patterns across FCI questions are heterogeneous and even incorrect answer choices provide meaningful information about students’ conceptions (27), we distinguished sub-types of “physics thinkers” based on their FCI answer choices. Specifically, we applied community detection to FCI answer distributions to identify sub-groups of similarly responding students and contrasted brain activity between groups to examine differential ways of thinking about the behavior of physical phenomena.

## RESULTS

### Physics problem solving engages visual motion, central executive, and default mode processes

FCI responses (mean accuracy = 61%, mean response time (RT) = 20.2s) were consistent with previous reports (27, 28) and significantly differed (p<0.001) from control responses (mean accuracy = 98%, mean RT = 15.8s), suggesting overall task compliance. Maps of FCI > Control blocks revealed activation across a fronto-temporo-parietal network, including the prefrontal cortex (PFC), left dorsal striatum, PPC, RSC, and dorsal posterior cingulate cortex, lateral occipitotemporal cortex (V5/MT+), and cerebellum (**Fig. 1a**; **Table S1**). To tease apart constituent neural processes, we analyzed sequential phases of the problem-solving process and observed multiple dissociable whole-brain networks linked with problem initiation (Phase I), question presentation (Phase II), and answer selection (Phase III). Phase I was associated with a similar activity pattern as the FCI > Control contrast, Phase II maps were characterized by right-emphasized dorsal posterior parietal and V5/MT+ engagement, and Phase III maps included medial anterior and posterior nodes of the default mode network (DMN; **Fig. 1b-d**; **Table S2**). These network transitions from fronto-temporo-parietal (Phase I) to dorsal attention (DAN; Phase II) followed by default mode cooperation (Phase III) points to the potentially important role V5-DMN-CEN interactions may have within physics reasoning processes. Meta-analytic functional decoding, which is a technique used to provide data-driven inferences about which mental functions are likely associated with specific brain activation patterns (see SI for more details), was performed on the resulting unthresholded z-statistic maps using Neurosynth (29), indicating that switching, default mode, motion perception, and reasoning processes may be important in physics problem solving (**Fig. 1** radar plots; **Table S3**).

**Fig. 1.**
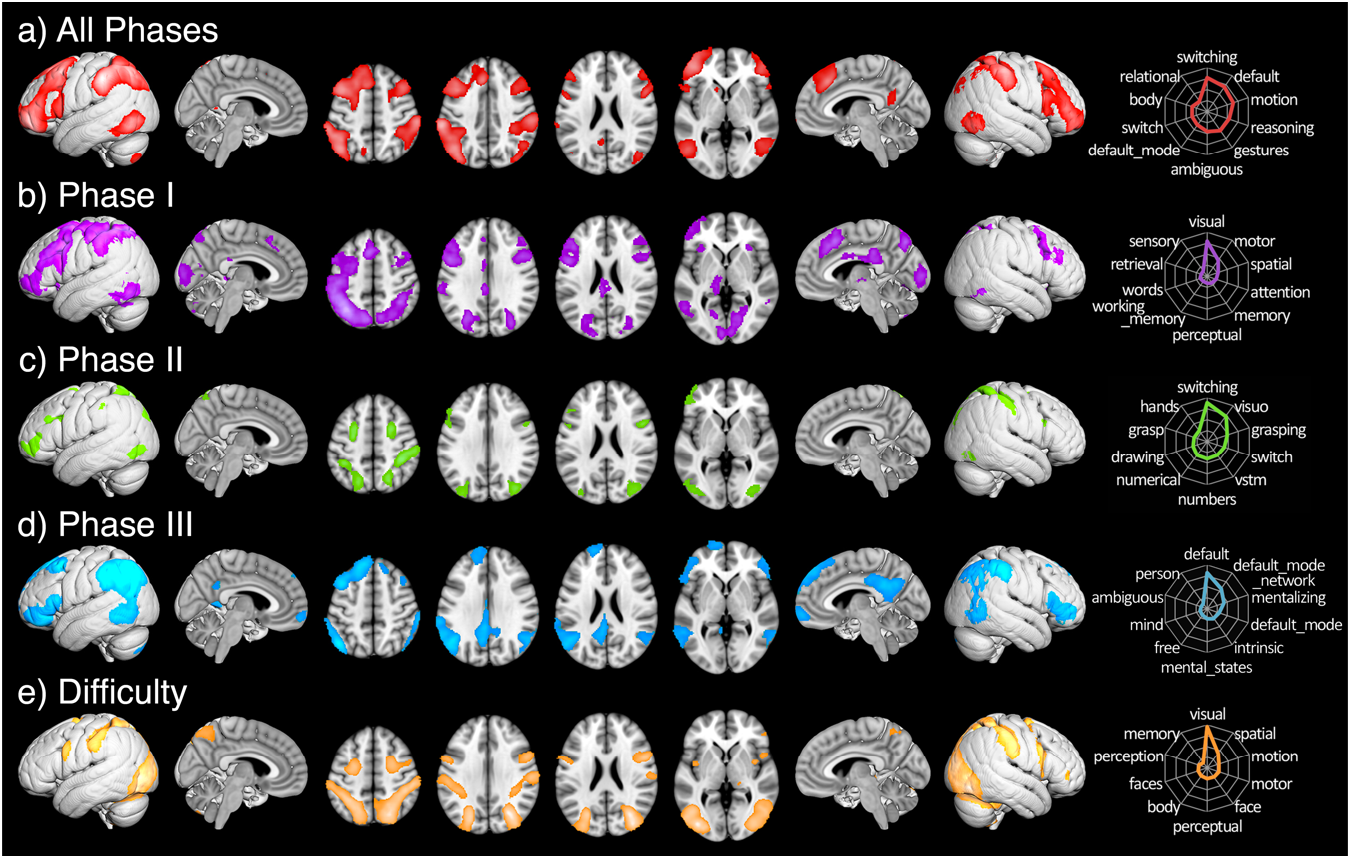
Activation of FCI > Control for a) problem solving across all phases, b-d) across each sequential problem phase, and e) parametric modulation across all phases by problem difficulty. Adjacent radar plots depict functional decoding results of the top ten weighted terms for each network.

Decoding sequential phases indicated problem initiation may reflect visuospatial attention, perceptual/motor, and memory retrieval; question presentation was associated with switching, visual short-term memory, and numbers, and answer selection was linked to DMN-related terms (e.g., unconstrained (free), mentalizing, and ambiguous), consistent with mental exploration of a solution. Next, to assess information exchange across GLM-identified regions during problem solving, we performed task-based functional connectivity (FC) analyses for three seeds centered on peaks of the overall FCI > Control map located in the left V5/MT+, the left dlPFC, and the RSC. Psychophysiological interaction (PPI) results (**Fig. 2**; **Table S4**) revealed greater physics problem solving-related coupling (relative to control conditions) of the left V5/MT+ with DAN brain areas, the left dlPFC with V5/MT+ and DMN areas, and the RSC with frontoparietal, DMN, and salience network (SN) regions. These outcomes suggest complex visual information may be carried through a dorsal stream to frontoparietal regions that direct CEN-DMN network exchanges during physics reasoning.

**Fig. 2.**
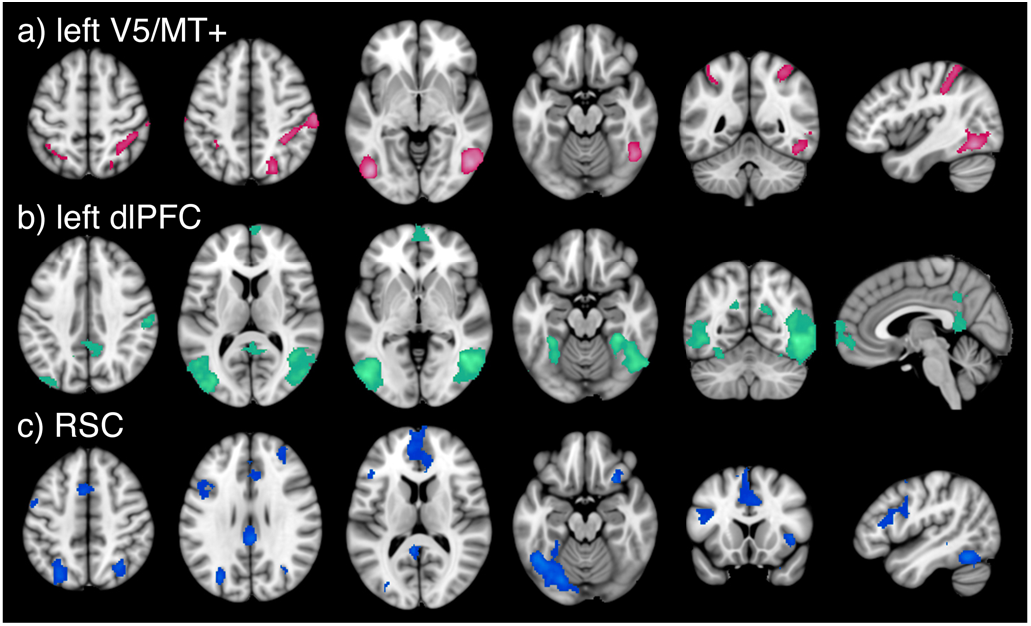
Whole-brain PPI task-based functional connectivity associated with FCI > Control for a) left V5/MT+, b) left dlPFC, and c) RSC seeds.

### Difficulty, but not accuracy and strategy, modulate brain activity during problem solving

To relate brain function to behavioral measures impacting student success, we tested our hypotheses that activity in meta-analytically derived ROIs (e.g., left dlPFC, left PPC, ACC, left hippocampus, and RSC) would be parametrically modulated by student-reported strategy and normative problem difficulty (30), but not answer accuracy. While no significant BOLD signal modulations were observed in these *a priori* ROIs, an exploratory whole-brain parametric modulation analysis revealed DAN and occipital activity were positively modulated by problem difficulty (**Fig. 1e**; **Table S5**). This indicates that the physics reasoning network is consistently activated regardless of whether or not a correct answer is achieved and does not reflect students’ perception of their reasoning strategy. Importantly, the most salient relation appears to be between degree of difficulty and engagement of brain regions linked with visuospatial perceptual, memory, and attentional processes, as assessed by functional decoding (**Fig. 1e** right). This may indicate mental states associated with memory, spatial or motion perception, and/or visualization may be especially engaged when problem difficulty is increased.

### Students demonstrate dissociable brain activity linked to knowledge fragmentation

We next performed module analysis (31) on students’ answer patterns to probe potential relationships between brain activity and students’ conceptual coherence (i.e., integration of physics knowledge; (22)) and to assess if distinct reasoning profiles were rooted in underlying functional brain differences. We analyzed answer distributions using a community detection algorithm (32) to parse student sub-groups who provided similar responses across FCI questions. Percent overlap was assessed between answers provided by each group and previously identified “conceptual modules” present in the FCI test ((31); **Table S6**). Conceptual modules are communities of incorrect FCI answer choices that are usually selected together. They represent students’ dissociable non-Newtonian (incorrect) notions about physical phenomena, some of which demonstrate a high degree of conceptual coherence, while others are more suggestive of a fragmented collection of physics ideas (21, 31, 33). The set of conceptual modules selected by a group (their reasoning profile) represents distinguishable arrangements of student’s (mis)interpretations and confusions about the physical world. Module analysis detected thirteen student groups across 107 students who answered similarly to each other during FCI problem solving (**Fig. 3a)**. Four groups had 10 or more members (i.e., normative groups). ANOVA indicated a significant difference in mean framewise displacement (FD) head motion between groups one or more of the groups (F(3, 178) = 8.213, p << 0.001). Post-hoc multiple comparison Turkey HSD tests indicated students in Group D showed to significantly greater head motion (p < 0.05). The three remaining the normative groups had no significant differences of in-scanner head motion and were thus selected for further analysis. The remaining three groups’ answer distributions were characterized based on prevalence of conceptual modules (**Fig. 3b)**. These groups, composed of 24, 17, and 10 students, were carried into group-level neuroimaging analyses to assess brain activity and connectivity differences during problem solving.

**Fig. 3.**
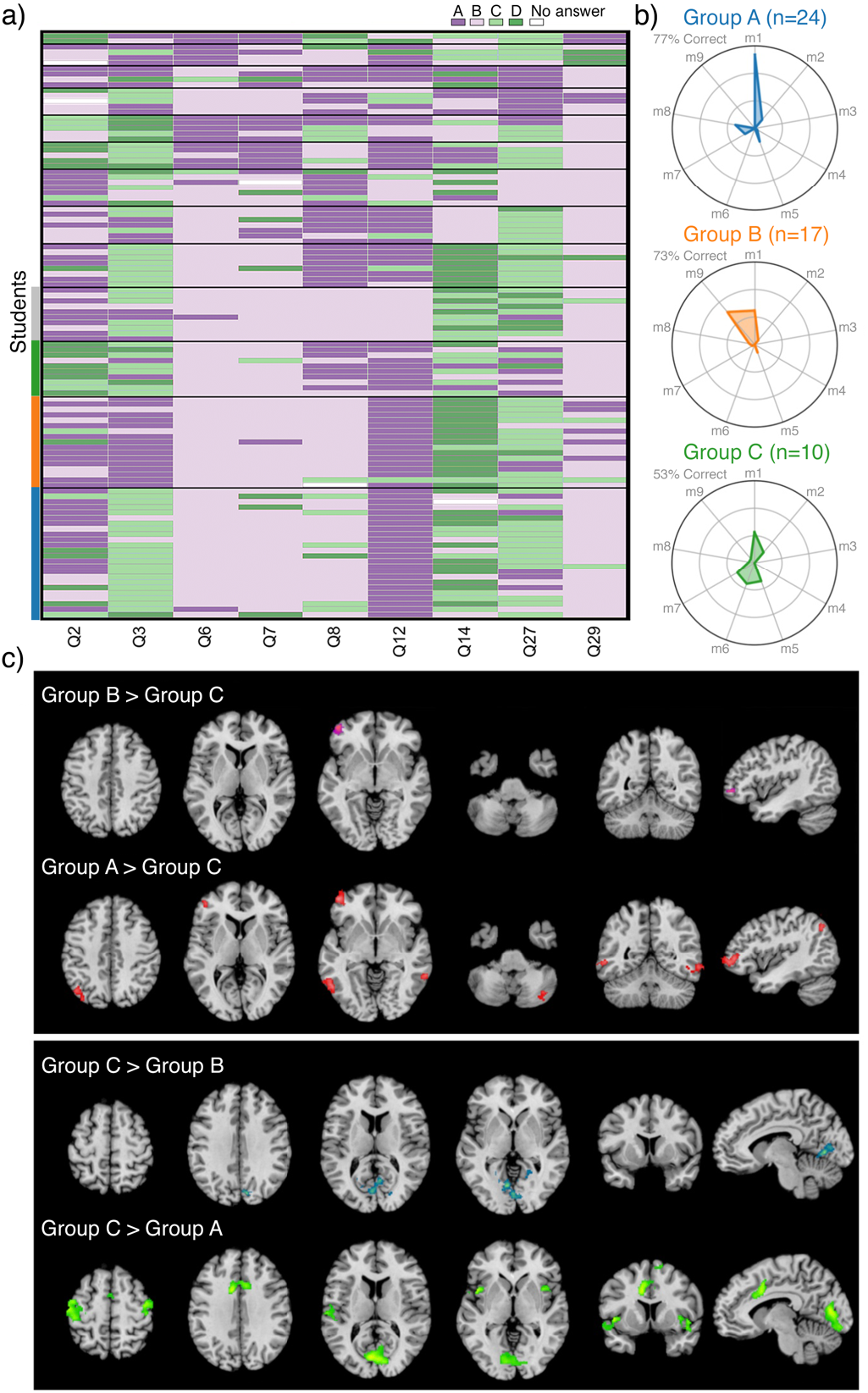
a) Module analysis of student responses across FCI answer distributions. Heat map colors represent student responses to multiple-choice FCI questions and black horizontal lines distinguish groups identified by community detection. b) Scaled within-group overlap of incorrect FCI responses across a nine previously measured physics conceptual models ((31); **Table S6**) for top three normative groups. c) Group differences in problem solving-related brain networks (FCI > Control, all phases) across the three normative groups. Increased activity is shown for Groups A and B relative to Group C (top) and Group C relative to Groups A and B (bottom). No significant differences were observed between Groups A and B.

Group A (n=24) achieved an accuracy rate of 77% across all FCI questions, indicative of being highly Newtonian thinkers (27). Of the non-Newtonian responses provided by this group, incorrect answers almost exclusively aligned with a common naïve physics idea known as the ‘*impetus force’* (*m1*, **Fig. 3b** top), which is the incorrect belief that moving objects experience a propelling force. Group B (n=17) achieved an accuracy rate of 73% across all FCI questions, which is also indicative of high Newtonian thinking. The reasoning profile for Group B (**Fig. 3b** middle) indicated that students gave incorrect answers by either falling victim to the *impetus force fallacy* (*m1*) or to another common, but less coherent set of physics conceptions that we term the ‘*confusion about gravitational action’ module* (*m9*). Group C (n=10) achieved an accuracy rate of 53% across all FCI questions, indicative of non-Newtonian thinking. The reasoning profile for Group C (**Fig. 3b** bottom) indicated that students’ incorrect answers were primarily associated with 5 conceptual modules that each occurred at relatively similar rates: the ‘*impetus force’ module* (*m1*), ‘*more force yields more result’ module* (*m2*), ‘*confusion relating speed and path’ module* (*m5*), ‘*sudden forces induce instantaneous path change’ module* (*m6*), and ‘an *object’s mass determines how it falls’ module* (*m7*).

We performed a whole-brain, one-way ANOVA to identify between-group differences in physics-related brain activity (FCI > Control, all phases). Omnibus results indicated that one or more sub-groups showed significantly different brain activity during problem solving. *Post hoc* tests were performed across each combination of group pairs (**Fig. 3c**; **Table S7**). Group A (vs. C) students demonstrated greater activity during problem solving in the left lateral orbitofrontal cortex (lOFC) as well as in the left inferior parietal lobule, bilateral V5/MT+, and right cerebellum. Group B (vs. C) students also exhibited greater activity in the left lOFC. Group C (vs. both A and B) students showed greater activity in the cuneus extending into the lingual gyri. Additionally, Group C students also showed increased activity relative to Group A in the caudal medial frontal gyrus, ACC, bilateral precentral and postcentral gyri along the precentral sulcus, bilateral anterior insular cortex (aIC), and left superior temporal gyrus. Overall, student who answered using more coherent physics conceptions, even if incorrect, showed increased reliance on a lOFC-V5/MT+ network, whereas students who held less consistent ideas involving multiple conceptual approaches showed increased primary visual and salience network activity. One possible interpretation of these differences may be that, in the absence of stable and coordinated physics conceptions, students engage relatively more visual search processes for salient problem features.

## DISCUSSION

Our fMRI results suggest that visualization, association, and mental exploration may be important mental processes that inform physics problem solving. When students solve physics problems they activate a network of bilateral dlPFC, left lOFC, PPC, RSC, and V5/MT+ areas, consistent with previous CEN-supported problem-solving findings across knowledge domains (5). Yet, V5/MT+ and RSC involvement with the CEN appear to be a feature of physics problem solving in particular. Both areas support visuospatial information processing (34), with V5/MT+ linked to imagining implied motion and maintaining motion information in working memory (35–37), and RSC supporting spatial cognition and episodic memory retrieval, especially when imagined scenes are mentally transformed between specific viewpoints (24). Thus, these regions may aid in the mental imagery of motion, as informed by remembered physical scenarios, and build internal representations of physical systems, which is considered an essential step in physics solution generation (38). Shifts in physics-related brain activity across problem phases indicate reliance on memory-linked associations. We find V5/MT+, CEN, DAN, and DMN transitions support sequential problem-solving phases. Notably, answer generation elicited concurrent DMN, lateral fronto-parietal, and V5/MT+ activity. Interestingly, while CEN-supported tasks often evoke DMN deactivations, this DMN-CEN coherence likely indicates reliance on episodic and semantic memory retrieval processes (39, 40) during physics cognition, a notion consistent with the constructivist theory of learning (41). Additionally, the PCC is functionally heterogeneous, connecting DMN and fronto-parietal networks, and serving as a possible hub across brain systems to direct attentional focus (42). Further, the FCI is differentiated from other fMRI tasks by its relatively long trials, requiring sustained cognition to generate answers. The DMN may thus be activated along with the CEN to allow for mental exploration necessary in solution derivation.

Problem solving-related brain activity was shown to differ based on how students think, not how correct they are. We found that students’ problem solving-related brain function cannot be categorized by simply considering their “*incorrect*” vs. “*correct*” answers. Rather, module analysis indicates variance in conceptual approach better characterizes brain differences, which in turn impacts success rate. An existing framework of learning conceptualizes physics cognition as relying on dual “knowledge structure” and “control structure” processes (22). Under this model, students apply executive functions to select or inhibit associational patterns that ground how they describe the physical world. Here, associational patterns, known as knowledge structures, are conceptualized as flexible, contextually-primed collections of linked knowledge elements called “resources” that students activate to scaffold reasoning. Ideally, students learn to activate stable associations between physical laws, enabling long deductive chains to be carried out during problem solving. However, when this does not occur, student’s non-Newtonian processes can vary: strongly associated yet inappropriate resources may stably activate across contexts, or more basic, axiomatic physical beliefs (e.g., intuitive notions such as *closer is stronger* or *more effort gives more result*; (21)) may form weak, unstable links that do not support ancillary deductive elaboration. These differences are described along an axis of “compilation” or memory chunking. Students without pre-compiled knowledge structures require additional cognitive resources to assemble associations during reasoning, whereas physics experts can access well-developed associational patterns that do not need to be actively assembled during problem solving.

We adopt this resources framework to interpret brain function with the goal of relating neuroimaging findings to educational knowledge and practice. Physics-related CEN and DAN activations were linked to varied cognitive terms consistent with the idea of a control structure, and DMN involvement during reasoning may reflect associational mappings within semantic or episodic memory circuits (39, 40). Thus, dlPFC-RSC FC may support the idea that control processes guide knowledge structure selection. Under this interpretation, reasoning sub-groups may be thought of as differentiated by knowledge structure use. Groups A and B applied predominantly Newtonian (i.e., compiled) thinking, but Group C was less consistent in their approach. Of the non-Newtonian modules activated, Group A consistently used an arguably concrete *impetus* model, Group B applied an *impetus* model while also expressing *confusion about gravitational action*, and Group C utilized multiple modules characterized by simple, vague, or confused ideas that differed across problems. We argue these groups can be described along a continuum of knowledge compilation, coherence, and robustness. Groups A and, to a lesser extent, B demonstrated stable, strongly associated knowledge structures, whereas Group C showed more labile associational patterns that were limited by problem context. In this manner, less coherent, more variable knowledge structures were associated with increased primary visual and SN activity, whereas pre-compiled, stable reasoning strategies more strongly activated lOFC and V5/MT+, areas implicated by physics thinking in the CEN. These findings suggest that chunked knowledge can reduce working memory demands, allowing for increased focus on other control structure aspects of problem solving (22). However, when students continually re-identify associational patterns across problems, they may rely more heavily on visually guided SN activity to select which problem features deserve their attention (43).

A fundamental goal of educational neuroscience is to bridge understanding of brain function with the insights, findings, and models of education research. Under a resources framework, our results suggest physics students struggle most when they do not understand how to choose appropriate and coherently chunked resources from long-term memory, thus relying on increased SN activity during problem solving. Learning obstacles also occur when students access compiled but non-physical conceptions during reasoning, allowing for increased CEN brain function linked to control processes. While the latter still represents a type of incorrect physics thinking, it more closely resembles the kind of cognition instructors aim to teach (22). As others have pointed out (44), it is a long path between brain imaging and the potential development of lesson plans, yet these insights may begin to inform aspects of physics classroom practice: instruction that explicitly attends to how students select, link, and reorganize resources may be critical in developing appropriately compiled knowledge to map back onto control processes (22). Learning physics is complex, yet a disproportionate focus is often placed on whether students answer questions correctly. Our results suggest the conceptual foundations of wrong answers are accompanied by functional brain differences during reasoning and can reveal much more about student’s ability to succeed than simple measures of accuracy. A focus on accuracy alone over-simplifies the complex processes engaged during physics reasoning. Instructors that leverage (rather than ignore or attempt to simply overwrite) students’ incorrect conceptions to facilitate conceptual change and transition existing resources about physical phenomena into stable and accessible knowledge structures may better serve students in connecting what they believe with what they predict.

In sum, we find the neural mechanisms underlying conceptual physics problem solving are characterized by integrated visual motion, central executive, attentional, and default mode brain systems, with solution generation relying on critical DMN-CEN engagement during reasoning. Furthermore, we explored whether measures of student success show underlying neurobiological bases, finding that students’ physics conceptions manifest as brain differences along an axis of relative knowledge fragmentation and robustness. Critically, accuracy alone did not predict brain function, but students achieved increased success when they made use of stable, strongly associated knowledge structures. We acknowledge that our results may be specific to the FCI questions used here, that additional or varied brain dynamics may be more relevant for different kinds of physics problem solving, and that sample sizes across Groups A, B, and C, are relatively small and uneven. Despite these concern, we are confident that our findings serve to deepen understanding into how students learn. Together, our results demonstrate associational and control processes operate in tandem to support physics problem solving and offer potential educational insight towards promoting student success.

## METHODS

### Participants

One hundred and seven healthy right-handed undergraduate students (age 18-25 years; 48 women) enrolled in introductory calculus-based physics at Florida International University (FIU) took part in this study. MRI data were acquired no more than two weeks after the end of the academic semester. Written informed consent was obtained in accordance with FIU Institutional Review Board approval.

### FCI Task

The Force Concept Inventory, a widely used (45) and reliable (46) test of conceptual understanding in Newtonian Physics (26), that includes a series of questions about physical scenarios was adapted for the MRI environment. FCI questions do not require mathematical calculation; rather they force students to choose between a correct answer and multiple common sense alternatives. The task included three phases: participants viewed a figure and descriptive text presenting a physical scenario (Phase I), a physics question was presented (Phase II), and participants viewed four possible answers and were instructed to choose the correct answer and mentally justify why their solution made the most sense (Phase III). Participants provided a self-paced button press to advance between phases and provide their final answer; a fixation cross was shown after answer selection before presentation of the next scenario. Question blocks were of maximum duration 45s and were followed by a fixation cross of minimum duration 10s. Control questions presented everyday physical scenarios and queried students on general reading comprehension instead of physics content. Control questions also included three phases (Control I, Control II, and Control III) to match the presentation of FCI questions. Post-scan debriefing included a paper-based questionnaire in which students rated the degree to which they had used “knowledge and reasoning” or had relied on a “gut feeling” to solve each FCI question.

### fMRI Acquisition and Pre-Processing

Functional images were acquired with an interleaved gradient-echo, echo planar imaging sequence (TR/TE = 2000/30ms, flip angle = 75°, FOV = 220×220mm, matrix size = 64×64, voxel dimensions = 3.4×3.4×3.4mm, 42 axial oblique slices). A T1-weighted series was acquired using a 3D fast spoiled gradient recall brain volume (FSPGR BRAVO) sequence with 186 contiguous sagittal slices (TI = 650ms, bandwidth = 25.0kHz, flip angle = 12°, FOV = 256×256mm, and slice thickness = 1.0mm). Pre-processing was performed using FSL (www.fmrib.ox.ac.uk/fsl) and AFNI (http://afni.nimh.nih.gov/afni) software libraries. Anatomical and functional images were skull stripped, the first five frames of each functional run were discarded, rigid-body motion correction was performed, functional images were high-pass filtered (110s), and a 12-degree-of-freedom affine transformation was applied to co-register the series with each structural volume. Non-linear resampling was applied to transform all images into MNI152 space and functional volumes were spatially smoothed using a 5mm Gaussian kernel. All motion-corrected non-registered 4D data underwent visual inspection and TRs associated with visually-identified motion artifacts were flagged for exclusion and their corresponding framewise displacement (FD) values were recorded. The minimum of the distribution of these artifact-linked FDs was used as a common scrubbing threshold across subjects during analyses. TRs with excessive motion (including one frame before and two frames after) were censored out during the GLM analysis if they met or exceeded a threshold of 0.35mm FD (47). Runs containing excessive motion (≥33% of within-block motion) were discarded from the analysis, resulting in the omission of three runs from two individuals. Six motion parameters (translations and rotations) were included as nuisance regressors in all analyses.

### General Linear Model Analyses

Stimulus timing files were created for each participant based on question phase onset/offset times. FCI and control questions were modeled as blocks from question onset to the onset of a concluding fixation cross triggered by answer selection. The contrast FCI > Control was modeled across full question duration; three additional GLM analyses were performed for the individual phases. Timing files were convolved with a hemodynamic response function and the first temporal derivatives of each convolved regressor were included to account for any offsets in peak BOLD response. General linear modeling for within-and between-subject analyses was performed in FSL using FEAT. Group-level activation maps for all contrasts were thresholded with a cluster defining threshold (CDT) of *P* < 0.001 and a cluster extent threshold (CET) of *P* < 0.05 (FWE corr).

### Task-Based Functional Connectivity Analysis

We tested for PPI associated with the FCI task across three seeds centered on peaks from the overall FCI > Control map located in the left V5/MT+, left dlPFC, and RSC. ROIs were transformed into native space and time series were extracted from unsmoothed data and included as regressors in separate within-subject PPI analyses performed on spatially smoothed 4D data sets. Design matrices for the within-subject PPI analyses contained regressors for the ROI time series, the condition difference vector modeling the differences between FCI and Control timing files, a vector representing the sum of the FCI and Control conditions, and the interaction between the task difference vector and ROI time series. The interaction term was calculated by zero-centering the task explanatory variable, and the mean of the ROI time series was set to zero. All task and interaction regressors, but not the ROI time series, were convolved with a Gamma-modeled hemodynamic response. PPI analyses were carried out separately for each ROI and resultant beta maps were averaged within-subject and carried into three separate group-level analyses. ROI-to-voxel task-based functional connectivity analyses were thresholded at a significance of *P* < 0.001 CDT, *P* < 0.05 CET (FWE corr).

### Brain-Behavior Correlates

Separate within-subject parametric modulation analyses were performed for accuracy, difficulty, and self-reported problem-solving strategy. Design matrices were identical to GLM analyses but included a single parametric modulator with the same FCI question timing but with a regressor height modeled by differences in the behavioral measures. Accuracy was modeled with regressor heights of 1, 0, or −1 corresponding to correct, no response, or incorrect answer provided. Difficulty was measured as a normative miss rate per FCI question, as measured externally (30). Problem-solving strategy was measured on a Likert scale by a post-scan questionnaire. If any parametric modulator had zero variance within a run (i.e., the student reported using an identical strategy for all questions) then the run was discarded to avoid rank deficiency in the design matrix. Resulting beta maps were then averaged across within-subject runs. Brain-behavior correlations were tested via two separate analyses: we extracted within-subject parametric modulator beta values within five hypothesis-driven ROIs and conducted one sample, two-sided t-tests on the beta distributions for significant variations from baseline (**Fig. S3**). Group-level analyses were also performed with whole-brain beta maps resulting from the parametric modulation GLMs to determine if significant network-level activity was present during problem solving associated with the behavioral measures.

### Student Response Profiles

Given evidence indicating student responses to the FCI provide insight into how students think about physics problems (31), we performed a module analysis of the observed FCI answer distributions to identify student response profiles. The data were treated as a bipartite matrix of Students × Responses. This bipartite matrix was computed and then projected into a weighted adjacency matrix of students, *A* = *MM*^*T*^, where *M* is the bipartite matrix. Each element in *A* represents the count of how many times one student agreed with any other student (values from 0 to 9, for 9 questions). Next, we performed nonparametric sparsification on *A* (48) to identify the backbone of the graph. Backboning identifies important links within a network and reduces the number of spurious links. A significance value was computed for each edge weight and the edge weights were thresholded at *P* < 0.01. We performed community detection (InfoMap R; (32)) on the backbone network to identify sub-groups of students who provided similar responses to the FCI prompts. We assessed the scaled within-group overlap of incorrect FCI responses across a set of nine previously measured physics modules consisting of jointly selected incorrect FCI response items ((31); **Table S6**). Each group’s relative conceptual module representation was scaled by group size to allow for comparisons across groups of different sizes. Alignment with conceptual modules indicates students draw on specific non-Newtonian physics conceptions. Finally, we tested for network differences across student groups. An omnibus test was conducted for the FCI > Control contrast as well as for the three whole-brain PPI maps. Significant F-test results were further interrogated with *post hoc* t-tests across groups. Maps were thresholded at *P* < 0.001 CDT, *P* < 0.05 CET (FWE corr).

### Data Availability

A GitHub repository was created at https://github.com/NBCLab/PhysicsLearning/tree/master/FCI to archive the data, code, and models for this study, including the e-Prime stimulus files, data analysis processing scripts, behavioral data, statistical brain images, and module analysis files.

## Supporting information

Supplemental Information (Revised)

## ACKNOWLEDGMENTS

Primary funding for this project was provided by NSF REAL DRL-1420627; additional support was provided by NSF 1631325, NIH R01 DA041353, NIH U01 DA041156, NSF CNS 1532061, NIH K01DA037819, NIH U54MD012393, and the FIU Graduate School Dissertation Year Fellowships. Thanks to Karina Falcone, Rosario Pintos Lobo, and Camila Uzcategui for their assistance with data collection and to the Department of Psychology of the University of Miami for providing access to their MRI scanner. Special thanks to the FIU undergraduate students who volunteered and participated in this project.

## AUTHOR CONTRIBUTIONS

**Designed research:** ARL, EB, SMP, MTS, RWL, JEB

**Performed research:** JEB, ERB, KLB, EIB, RO, AN

**Contributed analysis tools:** JEB, MCR, TS, KLB, EB

**Analyzed data:** JEB, MCR, TS, EB

**All authors contributed to the interpretation of the results and writing the manuscript.**

The authors declare no conflict of interest.

