## Supplemental Information (Revised) for "Brain activity links performance in science reasoning with conceptual approach"

**This PDF file includes:**

Supplementary text
Figs. S1 to S7
Tables S1 to S8
References for SI reference citations

### Supplementary Information Text

### SI Materials and Methods

**Participants.** One hundred and seven healthy right-handed participants who took part in this study were undergraduate students enrolled in introductory calculus-based physics at Florida International University (FIU) in Miami, Florida (age 18-25 years; mean: 20.2, SD: 1.39; 48 women). FIU is the fourth largest university in the US and is formally designated as both a Hispanic-Serving and Minority-Serving Institution (HSI and MSI, respectively). FIU is deeply committed to improving STEM education and ranks second in the US in number of STEM degrees awarded to Latinos and Latinas. Study participants were selected from a large set of applicants (N=496, from 22 different physics course sections) who responded to in-class recruitment announcements made at the beginning of the academic semester.

Participants were free of cognitive impairments, neurological and psychiatric conditions, did not use psychotropic medications (i.e. stimulants, anti-anxiety/anti-depressants, recreational drugs), and had never previously completed a university-level physics course. Of the 107 individuals who underwent post-instruction MRI scanning, 11 were college freshmen, 51 were sophomores, 32 were juniors, and 13 were seniors. The introductory physics course emphasized problem-solving skill development and covered topics in Newtonian mechanics, including motion along straight lines and in two and three dimensions, Newton's laws of motion, work and energy, momentum and collisions, and rotational dynamics. MRI scans commenced immediately after the completion of the physics courses final exam and concluded no more than two weeks after the end of the academic semester. Written informed consent was provided prior to participating in the study in accordance with FIU Institutional Review Board approval and students received monetary compensation for their time.

**FCI Task.** All tasks were programmed for presentation for the MRI environment using E-Prime (Psychology Software Tools, Inc., Pittsburgh, PA). Stimuli were projected onto a screen placed at the back of the MRI scanner and students viewed questions through a mirror view screen mounted to the head coil. Participants were given a fiber optic keypad to hold in their right hand with which to respond to each question. To probe the neural mechanisms underlying conceptual physics reasoning, we developed a scanner-adapted version of the *Force Concept Inventory* (FCI; (1)). FCI problems present physical scenarios involving objects at rest or in motion. Solution derivation requires extracting meaningful and relevant information about a scene, then appropriately applying physical laws to infer causal motion or interactions between forces and objects. The FCI is typically administered as an in-class multiple-choice exam consisting of questions about intuitive, every-day scenarios. We adapted and modified parts of the paper-based FCI exam for in-scanner display to accommodate presentation and timing requirements inherent to the MRI environment. All adaptions were made to remain as true as possible to the original in-class exam, with no changes fundamentally altering any physics-related content being tested. Original FCI question text was edited for brevity, placement of visual features was standardized across questions, and items were presented in a pseudo-randomized order. To encourage participant compliance and avoid fatigue or excessive head motion, we presented a reduced exam composed of 9 items from the original test. Included items (FCI 2, 3, 6, 7, 8, 12, 14, 27, and 29) probed student understanding of Newton's 1st and 2nd laws of motion. These questions were selected to span multiple difficulty levels (34.6% to 73.6% correct rate; (2)) and because their incorrect answer options probe a diversity of non-Newtonian conceptions about physics. Additionally, technological constraints associated with the four-button MRI-compatible keypad required that we eliminate one answer option from each of the originally five-answer choice FCI items. We removed the least commonly selected answer chosen by students across all ability levels, as reported in the item response curves of (2). These answer options were 2E, 3D, 6D, 7D, 8C, 12A, 14E, 27E, and 29C, and in-scanner FCI answer options appropriately were reordered.

In-scanner FCI and control questions were presented in a self-paced, three-phase sequence of view screens, emulating the flow of information in the original FCI exam (**Fig. S1a**). In the first problem initiation phase students viewed paired text and figure description a physical scenario (Phase I). Text was displayed on the top left portion of the view screen and did not exceed three sentences in length; the figure appeared at the top right portion of the view screen and depicted visual information necessary for answer making (e.g., kinematic trajectories or the spatial configuration of key features.) Students were instructed to press a keypad when they had completely read all text and felt they understood the physical scene. The button press triggered the start of the second question presentation phase (Phase II), which added a single, left-justified sentence to the middle portion of the view screen asking the student a physics question about the scenario. The student was instructed to press the keypad after fully reading the question in order to initiate the third and final answer selection phase (Phase III) wherein four possible answer choices, labeled A through D, were revealed at the bottom left of the view screen. Students were instructed to choose the correct answer and to explicitly mentally justify why the answer they selected made the most sense to them. Upon answer selection, all information on the view screen was replaced with a central fixation cross of variable duration. Question blocks were a maximum duration of 45s and were followed by a fixation cross of minimum duration 10s. Variable response times per block resulted in randomized interstimulus intervals between questions.

FCI questions were interleaved with control questions that did not require physics reasoning or problem solving, constituting a high-level baseline comparison (**Fig. S1b**). Control questions displayed text and figure depictions of everyday physical scenarios and tested students on general reading comprehension and/or shape discrimination instead of physics content.

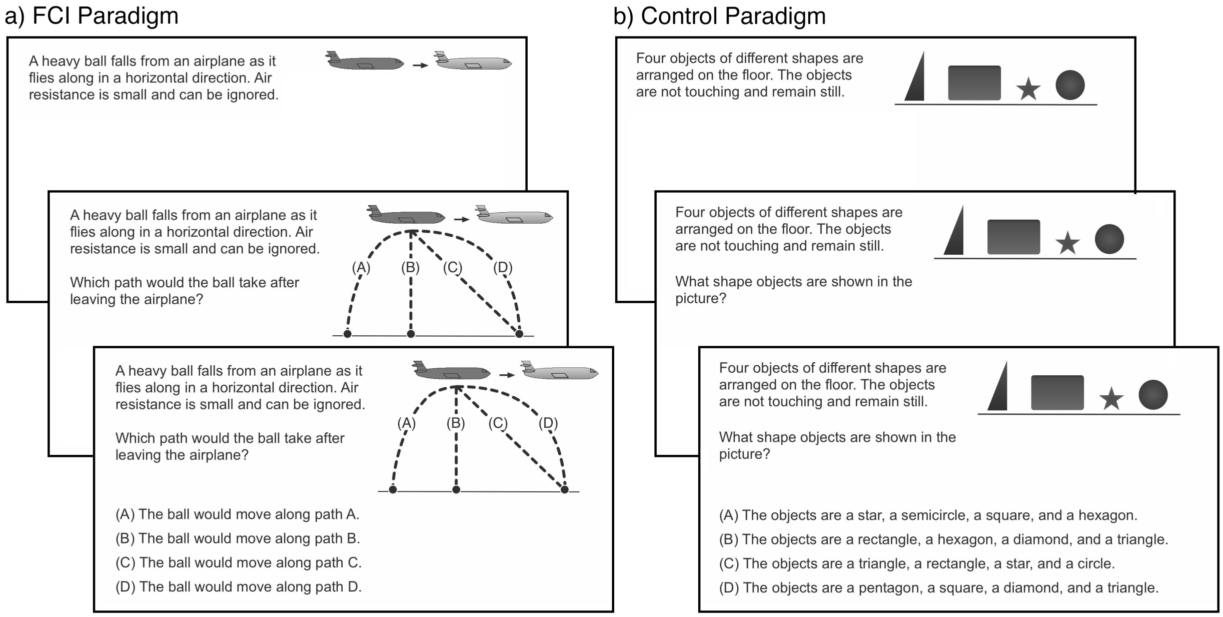


**Fig. S1.** Three-phase sequential progression of an exemplar in-scanner (a) Force Concept Inventory (FCI) question and (b) Control question. Additional images of FCI and Control questions can be viewed on GitHub (https://github.com/NBCLab/PhysicsLearning/tree/master/FCI/eprime-stimulus-files/fci-scanner/Slides).

Control items shared visual and linguistic characteristics to the FCI questions, containing words typically used in introductory Newtonian mechanics as well as visual presentation and self-paced timing paralleling that of the FCI problems. Text complexity for FCI and Control questions was measured using the Educational Testing Service's *TextEvaluator* tool (https://textevaluator.ets.org/textevaluator/) and no significant differences in linguistic complexity were present between conditions (total words per question: FCI = 31.4, Control = 31.3; Average words per sentence: FCI = 11.1, Control = 9.8; Syntactic complexity: FCI = 33.2, Control = 32.7; Academic Vocabulary: FCI = 22.7, Control = 32.6; Word Unfamiliarity: FCI = 36.7, Control = 32.3; Lexical Cohesion: FCI = 56.3, Control = 53.4; $p<0.05$).

**fMRI Data Acquisition and Pre-Processing.** Functional images were acquired on a GE 3T Healthcare Discovery 750W scanner with an interleaved gradient-echo, echo planar imaging (EPI) sequence (TR/TE = 2000/30ms, flip angle = 75°, field of view = 220x220mm, matrix size = 64x64, voxel dimensions = 3.4×3.4×3.4mm, 42 axial oblique slices, 172 volumes/run × 3 runs). A T1-weighted series was acquired using a 3D fast spoiled gradient recall brain volume (FSPGR BRAVO) sequence with 186 contiguous sagittal slices (TI = 650ms, bandwidth = 25.0kHz, flip angle = 12°, FOV = 256x256mm, and slice thickness = 1.0mm). Pre-processing was performed using tools from the FSL (www.fmrib.ox.ac.uk/fsl) and AFNI (http://afni.nimh.nih.gov/afni) software libraries. To allow for image intensity stabilization, the first five frames of each functional run were discarded. All functional and structural images were aligned to a common stereotactic origin and spatial orientation to match that of the MNI152 template. Rigid-body motion correction was performed on functional runs by aligning all images in each run to the middle volume. Anatomical and functional images were skull stripped, functional images were high-pass filtered (110s), and a 12-degree-of-freedom affine transformation was applied to co-register the series with each participant's structural volume. Non-linear resampling was applied to transform all images into MNI152 2mm space and functional volumes were spatially smoothed using a 5mm Gaussian kernel. Additionally, all motion-corrected non-registered 4D data underwent visual inspection and TRs associated with visually identified motion artifacts were flagged for exclusion in further analysis and their corresponding FD values were recorded. The minimum of the distribution of these artifact-linked FDs was used for a common scrubbing threshold across subjects during general linear modeling.

**FCI Problem Difficulty and Post-Scan Strategy Questionnaire.** We wished to assess any neural processes that were parametrically more associated with using formal deduction or physical intuition when solving physics problems, two separate processes that we refer to collectively as students' problem solving strategy. We additionally wished to assess any neural activity that was parametrically more activated when problem difficulty was increased. Problem solving strategy was measured as a self-reported measure assessed immediately after scan completion. Students were given a post-scan, written questionnaire depicting each in-scanner FCI question with the statements "*I used knowledge and reasoning to arrive at my answer*" and "*I relied on a 'gut feeling' to arrive at my answer*." Students rated their agreement/disagreement with each statement for each FCI question on a 5-point Likert scale. Normative question difficulty was measured as the percent of students who answered incorrectly on FCI questions from a dataset of more than 4,500 student responses to the FCI administered at Harvard University, Mississippi State University, and Rice University and reported by Morris (2).

**General Linear Model and Parametric Modulator Analyses.** Stimulus timing files were created for each participant based on question and phase onset/offset times. FCI and control questions were modeled as blocks from question onset to the onset of a concluding fixation cross triggered by answer selection. The contrast FCI > Control was modeled across full question duration; three additional GLM analyses were performed for the individual phases. Timing files were convolved with a Gamma-modeled hemodynamic response function and the first temporal derivative of each convolved regressor was included in analyses to account for any offsets in peak BOLD response. TRs with excessive motion were scrubbed if they met or exceeded a threshold of 0.35mm FD, as determined by the visual inspection of minimally preprocessed data. This was accomplished by including separate, single-TR regressors in the design matrices of subject-level GLM analyses if excessive motion was observed, thereby censoring out high motion volumes during parameter estimation. To account for temporal blurring of BOLD signal, one frame before and two frames after each TR with excessive motion were additionally scrubbed from the dataset (3). Runs containing excessive motion (≥33% of within-block motion, as modeled across all questions within a run) were discarded from the analysis, resulting in the omission of three runs from two individuals. Six motion parameters (translations and rotations) were included as nuisance regressors in all analyses. Parametric modulator analyses contained identical design matrices to those of the FCI > Control (all phases) subject-level analyses but included a parametric modulator regressor wherein question duration was modeled by student-specific FCI > Control response times and regressor heights were modulated by question-specific accuracy, self-reported strategy (as assessed by post-scan strategy questionnaires), and question difficulty. Accuracy and problem solving strategy were subject-specific measures and question difficulty was modeled externally as a normative metric of how challenging (% incorrect) each FCI question generally is for introductory physics students, as measured across a large body (>4,500) of university students who had taken the exam (2). General linear modeling for within- and between-subject analyses was performed in FSL using FEAT. Group-level activation maps for the contrasts FCI > Control, Phase I > Control I, Phase II > Control II, and Phase III > Control III, and for the whole-brain parametric modulator analyses were thresholded using a cluster defining threshold (CDT) of $P < 0.001$ and a cluster extent threshold (CET) of $P < 0.05$ (FWE corrected). Meta-analytic functional decoding for the FCI > Control, Phase I > Control I, Phase II > Control II, Phase II > Control III, and Difficulty Modulator contrasts was performed on the resulting unthresholded z-statistic maps with a 200-topic GC-LDA (4) topic model trained on the Neurosynth database, a methodology that is explained in greater detail below.

**Meta-analytic Functional Decoding Using Neurosynth.** Understanding the relationship between a specific pattern of brain activity and which underlying cognitive states it is associated with is one of the fundamental goals of neuroimaging research (5). Typically, as in this study, neuroimaging experiments are performed to test the forward inference problem: given a specific mental process, what is the observed pattern of brain activity? It is also desirable (and frequently the goal of discussion sections) to examine the broader cognitive processes that are linked to the observed pattern of brain activity. Yet, the reverse problem (given a pattern of brain activity, which cognitive or psychological processes likely produced it?) is significantly more complicated to answer. An example illustrating this challenge is the case of the amygdala – experiments in which individuals were presented with fearful stimuli often show associated amygdala activity (6), yet due to the many-to-many nature of how mental states map onto spatially localized brain function, the reverse inference need not be true. That is, an observation of activity in the amygdala does not necessarily indicate the neural process was due to an underlying fearful response. Indeed, claiming the presence of one or more underlying mental states based on the results of a limited set of experiments is a logical fallacy. Fortunately, the reverse question can still be answered in a way that avoids this error: one can compute the probabilistic correspondence between a specific result and a wider (e.g., representative of the broader neuroimaging literature) set of mental states and associated activation patterns (5). In this way researchers can formally assess the likelihood that any given mental state is associated with a particular activity pattern.

Probabilistically leveraging the wider neuroimaging literature to guide the interpretation of experimental results is an arguably more objective approach than speculating about the meaning of results based on only a limited set of citations brought into the discussion. Unsurprisingly, many researchers have begun to adopt this technique and increasingly sophisticated probabilistic tools and frameworks for 'decoding' brain activity have been developed (4, 7, 8). Neurosynth (http://neurosynth.org; 7) is a large database (>14,000 neuroimaging studies) of mappings between neural states and cognitive processes that can be used to perform quantitative reverse inference. Neurosynth uses automated text mining algorithms to access published neuroimaging studies and extract meaningful words (or 'terms') from the paper's abstract along with peak activation coordinate results as reported in the paper's tables. Terms are thought to represent cognitive and/or psychological processes that are linked to the activation coordinates. In this paper, we make use of a Generalized Correspondence Latent Dirichlet Allocation (GC-LDA) machine learning algorithm (Rubin, 2016) which was trained on the Neurosynth database to determine likely sets of underlying mental functions that are associated with our observed FCI-related brain activity. GC-LDA is a data reduction technique that was used to detect a set of latent and semantically coherent 'topics' that are present in the Neurosynth database. Each topic consists of paired sets of spatial activation coordinates and semantically linked terms. Thus, topics form highly interpretable collections of functional brain regions linked with mental processes that can then be used to decode whole-brain activation patterns (8). In this study we fed in the unthresholded z-statistic brain activation FCI-related maps into the GC-LDA topic model using 200 distinct topics to determine a rank ordered lists of terms that, according the latent topics present in the Neurosynth database, are most associated with the input pattern of brain activity observed during the FCI. We did this to form a data-driven text-based representation of each observed activation pattern so as to objectively guide our interpretation of results.

**Definitions of *A Priori* Regions of Interest and PPI Seeds.** Five *a priori* regions of interest (ROIs) were selected for inspection of potential physics problem solving-related brain activity correlations with problem solving strategy, accuracy, and difficulty. ROIs were meta-analytically defined to include areas associated with *problem-solving* (e.g., left dorsolateral prefrontal cortex (dlPFC), ACC, left posterior parietal cortex (PPC) (9)) and *episodic and spatial memory* (e.g., left hippocampus and retrosplenial cortex (RSC); (10, 11)). A recent meta-analysis on problem-solving revealed the left dlPFC, the left PPC, and the ACC as critically involved in problem solving involving mathematical, visual, or verbal stimuli (9). Centroid meta-analytic coordinates from these regions were used as seeds in the present analysis. In addition, to investigate to putative connection between memory-related processes during physics problem-solving and behavioral measures, we selected two functionally-relevant ROIs from memory neuroimaging literature in the left hippocampus and RSC to explore potential involvement long-term and episodic memory retrieval plays within physics reasoning. These two regions were chosen to investigate the role long-term memory and/or autobiographical memory, especially when involving spatial thinking, may have on behavioral measures during physics problem-solving. For the hippocampal seed, a region was chosen the left middle hippocampus based on connectivity-based parcellation findings suggesting this region is particularly involved in declarative memory (10). The RSC seed was drawn from peak coordinates from meta-analytic results on the neural correlates on autobiographical memory (11). In that study, the RSC was identified as a region simultaneously present within a core autobiographical memory network, as well as particularly involved in memory retrieval of events characterized by spatial context and visuospatial processing. The five ROIs were modeled as 10mm spherical seeds (**Fig. S2a**).

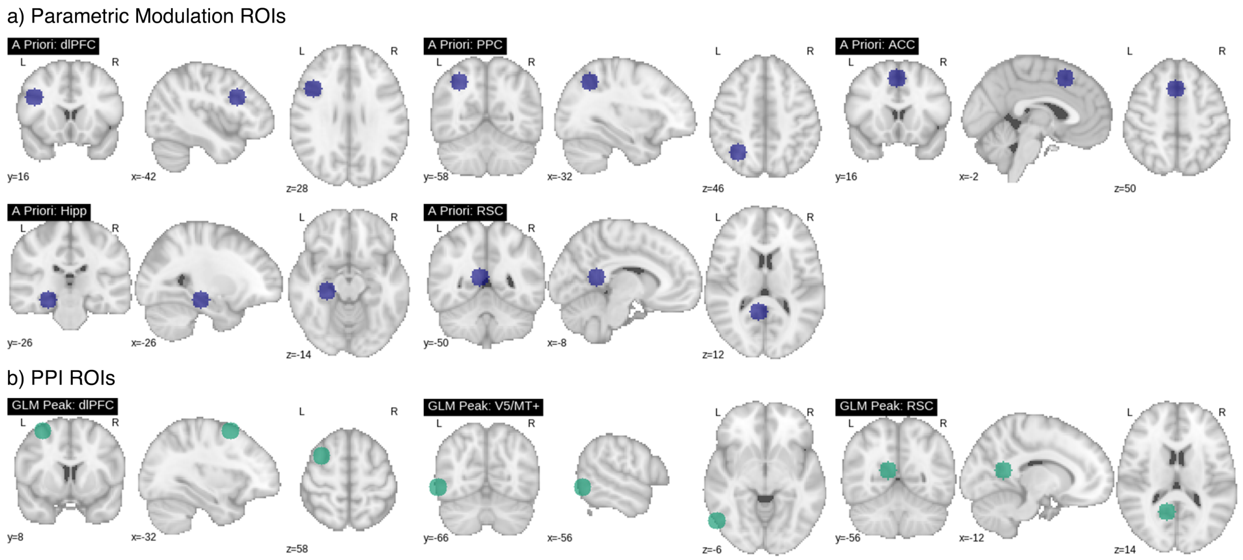


**Fig. S2.** a) The hypothesis-driven ROIs (blue) selected from coordinate results of problem solving (dlPFC, PPC, and ACC; (9)) and episodic, spatial, and declarative memory (hippocampus and RSC; (10, 11)) meta-analyses. b) The seeds selected for further exploration of task-based functional connectivity via psychophysiological interaction analysis (PPI; green). These regions were derived from peak group-level results from the FCI > Control (all phases) contrast.

Solving physics problems relies on deduction and knowledge recall; thus, we hypothesized that the fMRI signal in the problem-solving and hippocampus ROIs would parametrically increase with problem difficulty and reasoning strategy. Specifically, we expected difficulty to modulate activity in ACC and dlPFC and reasoning strategy to modulate activity in ACC, dlPFC, PPC, and hippocampus. If students reported using physical intuition (i.e., they answered via a *"gut feeling"*), we expected a positive parametric effect in RSC, an area linked to visualization and memory of autobiographical experiences (12). Additionally, due to the influence of strongly held yet non-physical conceptions on confidence bias in introductory Newtonian mechanics (13), we did not expect accuracy-related parametric effects to be present in any ROI.

Separate, *post hoc* analyses were conducted using a psychophysiological (PPI) approach to explore task-based functional connectivity between brain regions implicated as supporting physics cognition as determined by General Linear Model analysis. The three ROIs used in the PPI analyses were centered on peak coordinates of the group-level overall FCI > Control contrast map in the left dlPFC, left V5/MT+, and RSC and were modeled as 10mm spheres (**Fig. S2b**).

**Description of Conceptual Modules and How They Were Computed.** Recent work has identified distinct communities of non-Newtonian FCI answer choices given by frequency of co-occurring student responses to the original FCI exam (14). These so-called "conceptual modules" represent dissociable incorrect physics conceptions that students commonly hold. Each of the nine identified modules have been interpreted based on the shared incorrect physics conceptions present within each community of answer choices (14). Similar communities of incorrect FCI answer choices have been identified and analogously interpreted in other investigations (1, 15). According to these characterizations, some communities show a high degree of conceptual coherence similar to what some education researchers have historically termed a physics "misconception" (that is, the collection of concepts that represent an integrated model of incorrect physics ideas students' typically apply across diverse contexts; (1, 16, 17)), while others are more consistent with a "knowledge-in-pieces" view of student physics thinking (14, 18) in which students sometimes reason through physics problems by drawing upon knowledge elements that are more fragmented or loosely connected (16, 17, 19). Students who rely on less coherently organized knowledge structures such as these tend to have a difficult time applying the same knowledge across contexts and situations (17).

Conceptual modules were identified (14) through a process of treating student FCI answer responses as a bipartite network represented as a Students X Responses matrix. This matrix was then multiplied by its transpose to project the bipartite network into a Responses X Responses matrix, which was weighted by the number of students choosing each answer pair. So for example, assuming three students choose answer A on FCI question 2, and two of these three students chose answer C on FCI question 3 with the third student choosing D on FCI question 3, then the answer projection of the bipartite network would be an edge with weight 2 between answers 2A and 3C and a edge with weight 1 between answers 2A and 3D. At this point, the answer projection network represents the network of responses, but it is too densely connected to analyze. Backboning or sparsifying the network is a process that aims to reduce the number of edges by retaining only the 'important' edges while preserving as many connected nodes as possible. In order to sparsify this network, the researchers used a locally adaptive non-parametric sparsification (LANS) algorithm (20) which works with non-parametric distributions. A community detection algorithm (InfoMap R; D. Edler and M. Rosvall, The MapEquation software package, available online at http://www.mapequation.org; (21)) was then applied to sparsified network to identify groups of responses that are more commonly connected together than to the rest of the network.

### SI Results

**GLM Results.** Center of mass activation coordinates for the FCI > Control contrast are shown in **Table S1**. Coordinate results for the contrasts FCI Phase I > Control Phase I, FCI Phase II > Control Phase II, and FCI Phase III > Control Phase III are shown in **Table S2**. Meta-analytic functional decoding results via NeuroSynth are shown in **Table S3**. **Table S4** displays the coordinate results for the PPI analyses for the dlPFC, RSC, and V5/MT+ seeds.

**Brain-Behavior Correlates.** Brain-behavior correlations were tested via two separate analyses. In the first analysis we extracted within-subject parametric modulator beta values within the five hypothesis-driven ROIs and conducted one sample t-tests on the beta distributions for significant variations from baseline. In this first analysis we tested the hypotheses that 1) fMRI signal in the ACC and the left dlPFC would parametrically increase with problem difficulty, that 2) signal in the ACC, dlPFC, PPC, and hippocampus ROIs would parametrically increase with reasoning strategy (i.e., they answered using *"knowledge and reasoning"*), that 3) fMRI signal in the RSC would parametrically increase if students reported using physical intuition (i.e., they answered via a *"gut feeling"*), and that 4) no accuracy-related parametric effects would be present in any ROI. No significant variations in BOLD signal from baselines were observed within the ROIs tested. Beta distributions across all ROIs are shown in **Fig. S3**.

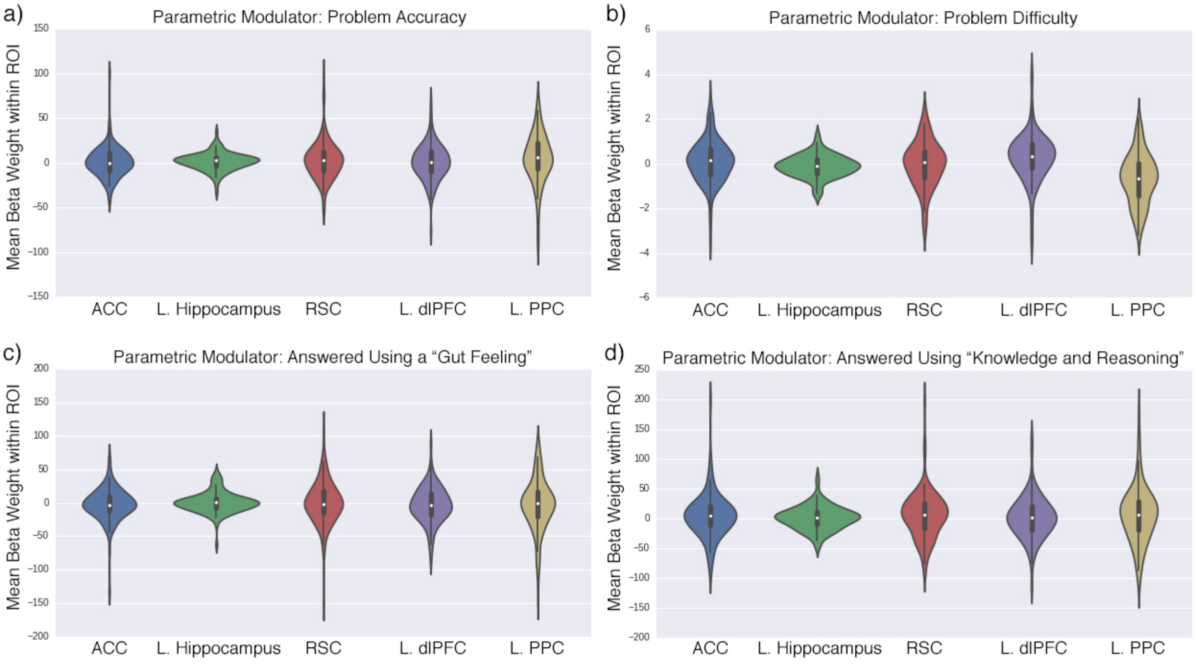


**Fig. S3.** Mean beta weight distributions for the four parametric modulator analyses within the *a priori* ROIs. a) Problem accuracy, b) difficulty, and c-d) strategy are shown.

In the second analysis, whole-brain beta maps resulting from the parametric modulation GLMs were averaged across groups to determine if significant network-level activity, outside that of any selected hypothesis driven ROIs, was present during problem solving associated with the behavioral measures.

Similar to the *a priori* ROI analyses, no significant results were observed for the Accuracy or strategy modulators, but significant whole-brain group-level modulation was linked with the difficulty regressor (**Table S5**). These results are described in detail in the main text.

**Normative Reasoning Groups.** Module analysis of students' answer patterns identified nine sub-groups. The resulting groups of responses were interpreted, much like in factor analysis, to identify common themes among the responses in the modules. The constituent FCI answer choices for each module and a summary of the themes are provided in **Table S6**.

We next sought to identify any differences in physics problem solving-related brain function associated with differences in conceptual approach by contrasting the brain activity of the student sub-groups. While all fMRI data had been scrubbed with a common framewise displacement (FD) threshold during preprocessing to eliminate visually identifiable artifacts, not all head movement artifacts are identifiable via visual inspection and even small motions can cause signal changes in fMRI data that can interfere with the interpretation of results (22). So, to avoid any potential motion-related confounds during group comparison of brain function, a one-way ANOVA of mean FD values across the four normative ($n \geq 10$) groups was conducted and a significant difference of in-scanner motion ($F(3, 178) = 8.213$, $p << 0.001$) was detected (mean FD: Group A = 0.072mm, Group B = 0.062mm, Group C = 0.073mm, and Group D = 0.092mm). *Post hoc* tests revealed a single normative group (Group D) showed significantly increased motion relative to all other normative groups ($p < 0.05$; **Fig. S4**), but no other differences in in-scanner motion existed.

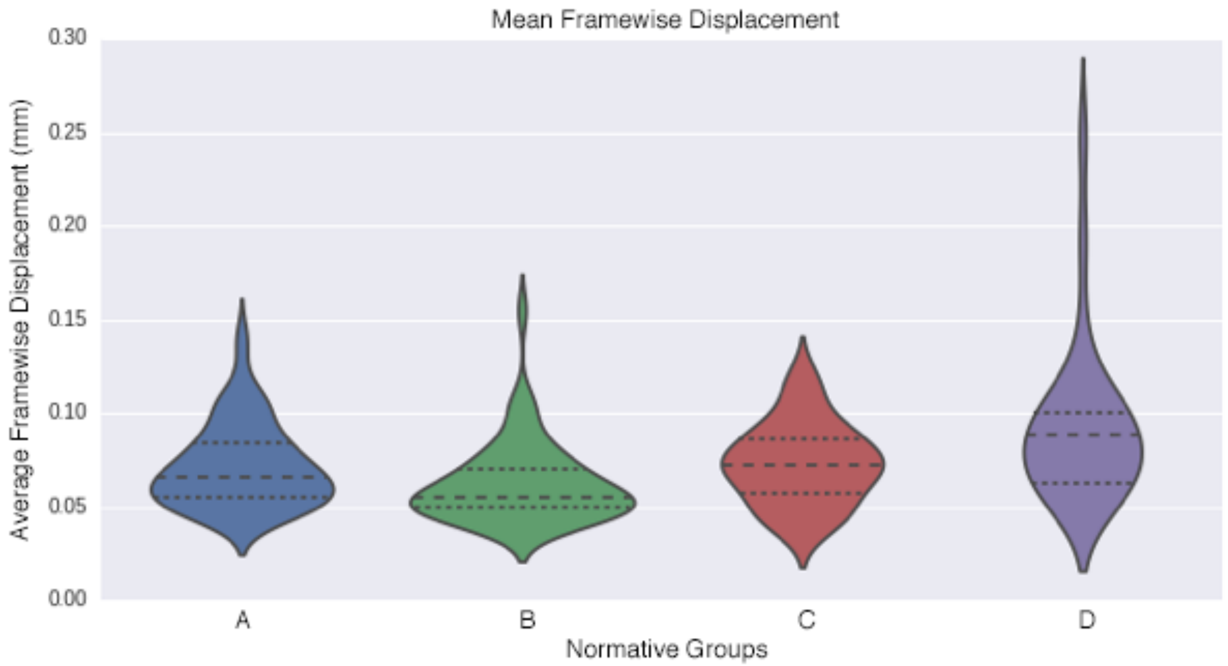


**Fig. S4.** Mean framewise displacement (FD; mm) for the four normative ($n \geq 10$) sub-groups identified by module analysis of FCI answer distributions. ANOVA indicated a significant difference in mean FD head motion between groups one or more of the groups ($p << 0.001$). Post-hoc multiple comparison Turkey HSD tests indicated students in Group D showed to significantly greater head motion ($p < 0.05$) relative to groups A-C, thus Group D was excluded from further analysis.

Thus, to avoid any potential confounds related to differences in head motion across normative groups, the high motion group was excluded from further analyses. Center of mass activation coordinates for the FCI > Control contrast of group differences between the final three normative sub-groups identified by module analysis are shown in **Table S7**.

**Response Times.** Average response times (RT) for FCI and control questions across all students were 20.2s and 15.7s respectively (FCI Phase I: 6.4s, Control Phase I: 6.8s; FCI Phase II: 5.1s, Control Phase II: 2.8s; FCI Phase III: 8.6s, Control Phase III: 6.1s; **Fig. S5**). Reasoning sub-group FCI and control RTs (across all phases) were 21.2s and 15.7s for Group A, 18.1s and 14.4s for Group B, and 20.1s and 16.8s for Group C (**Fig. S6**). We conducted statistical comparisons to determine if differences in RT were present across problem phases, conditions, and reasoning sub-groups.

In a comparison across all 107 student responses, significant RT differences were detected between FCI and Control conditions across full question blocks (all phases; $p << 0.001$). A two-way mixed effects ANOVA: condition (FCI, Control) x Phase (Phase I, Phase II, Phase III) was also conducted to compare the main effects of phase and condition and the interaction between phase and condition on RT. A significant two-way interaction between phase and condition was detected ($F(2,535) = 62.860$, $p << 0.001$), indicating RTs differed across conditions and phases. Turkey HSD *post hoc* tests were conducted on the family of six estimates and all but two pairwise comparisons were significant (see **Table S8** for a summary these multiple comparisons.) No significant RT differences were detected between conditions at Phase I, or between Phase III FCI RT and Phase I Control RT. All other comparisons were significant at $p < 0.001$ except for the Phase I Control RT - Phase III Control RT contrast, which was significant at $p < 0.05$. These results indicate students spent significantly more time answering FCI questions as compared to Control questions, except within Phase I. Students also spent significantly more time in Phases III, I, and II, respectively across both conditions.

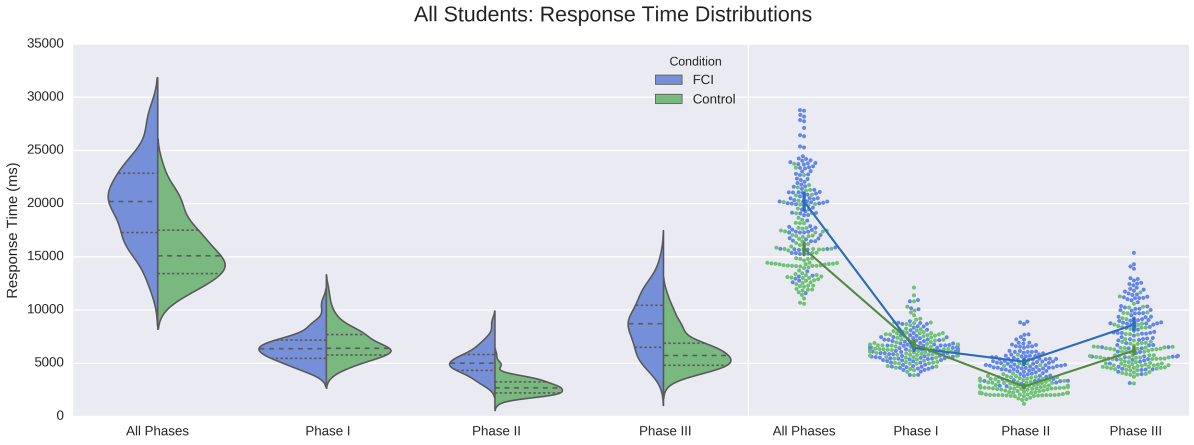


**Fig. S5.** Mean response times (RT; ms) across all students for the FCI and Control condition (all phases) as well as each sequential phase. Kernel density estimates of RT distributions are plotted on the left and the interaction across conditions is provided on the right, with 95% confidence intervals shown as vertical error bars. Students spent significantly more time answering FCI questions compared to Control questions, except within Phase I, and significantly more time in both conditions in Phase III, Phase I, and Phase II, respectively ($p << 0.001$).

We also tested if response times were correlated with mean FCI performance or strategy (e.g., student's self-reported statements on how much they relied on "knowledge and reasoning" and/or a "gut feeling" to answer the FCI questions). Pearson correlation values of response times with these measures were computed for each Phase as well as across full FCI questions. Associated p-values were assessed and correction for multiple comparisons was carried out to determine significance using the Bonferroni method. No significant correlations were observed between response times and FCI strategy (FCI Phase I: $p_{\text{knowledge and reasoning}} = 0.9556$ and $p_{\text{gut feeling}} = 0.2872$; FCI Phase II: $p_{\text{knowledge and reasoning}} = 0.\ 1512$ and $p_{\text{gut feeling}} = 0.\ 1458$; FCI Phase III: $p_{\text{knowledge and reasoning}} = 0.\ 1841$ and $p_{\text{gut feeling}} = 0.\ 1704$; FCI All Phases: $p_{\text{knowledge and reasoning}} = 0.\ 1729$ and $p_{\text{gut feeling}} = 0.\ 0702$). Response time was significantly correlated with accuracy for Phases I and II, but not for Phase III or across all Phases (FCI Phase I: $p_{\text{accuracy}} < 0.05/12$; FCI Phase II: $p_{\text{accuracy}} < 0.05/12$; FCI Phase III: $p_{\text{accuracy}} = 0.8257$; FCI All Phases: $p_{\text{accuracy}} = 0.0227$). **Fig. S6** shows scatter plots for accuracy and response time across these Phases.

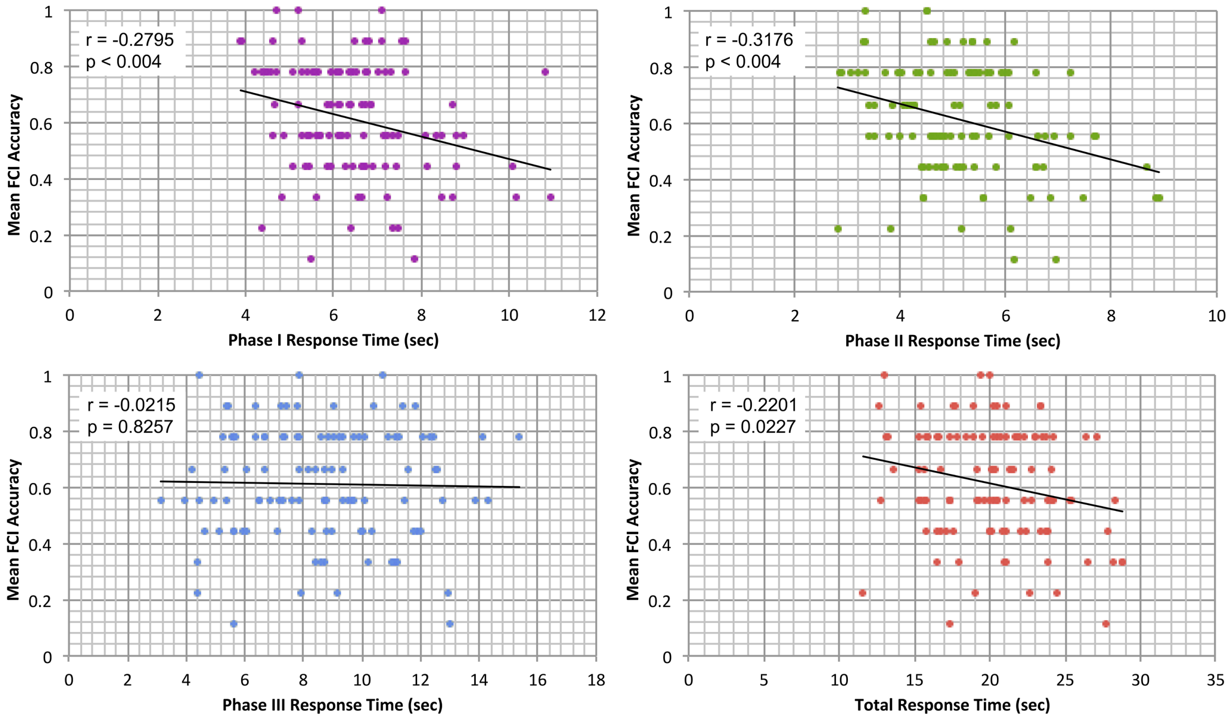


**Fig. S6.** Scatter plots of FCI response times with task performance are shown across each Phase and across the full FCI task. Pearson correlations are provided for each comparison along with Bonferoni corrected p values. Response time was significant correlated with accuracy for Phase I and Phase II but not for Phase III or overall across FCI questions.

Additionally, to investigate potential RT differences across normative reasoning groups, we conducted one-way ANOVAs testing within-condition mean RT across reasoning sub-groups. A significant difference in RT across sub-groups was detected for the FCI condition ($F(2,48) = 4.315$, $p < 0.05$) but not for the Control condition RTs ($F(2,48) = 2.269$, $p = .114$). However, turkey HSD *post hoc* tests revealed no significant pairwise FCI RT differences between normative groups after correcting for multiple comparisons. A two-way mixed effects ANOVA: condition (FCI, Control) x Group (Group A, Group B, Group C) was also conducted to compare the main effects of group and condition and the interaction between group and condition on RT. The effect of group was significant, yielding an F ratio of $F(2,48) = 3.8139$, $p < 0.05$. However, as before, *post hoc* multiple comparison Turkey HSD tests indicated no significant pairwise differences between groups ($p < 0.05$, adjusted for multiple comparison using the Holm method.) The effect of condition yielded an F ratio of $F(1,48) = 100.5341$, $p << 0.001$, indicating a significant difference in RT between FCI and Control conditions. The interaction effect between condition and group was not significant, $F(2,48) = 2.1661$, $p = 0.1257$.

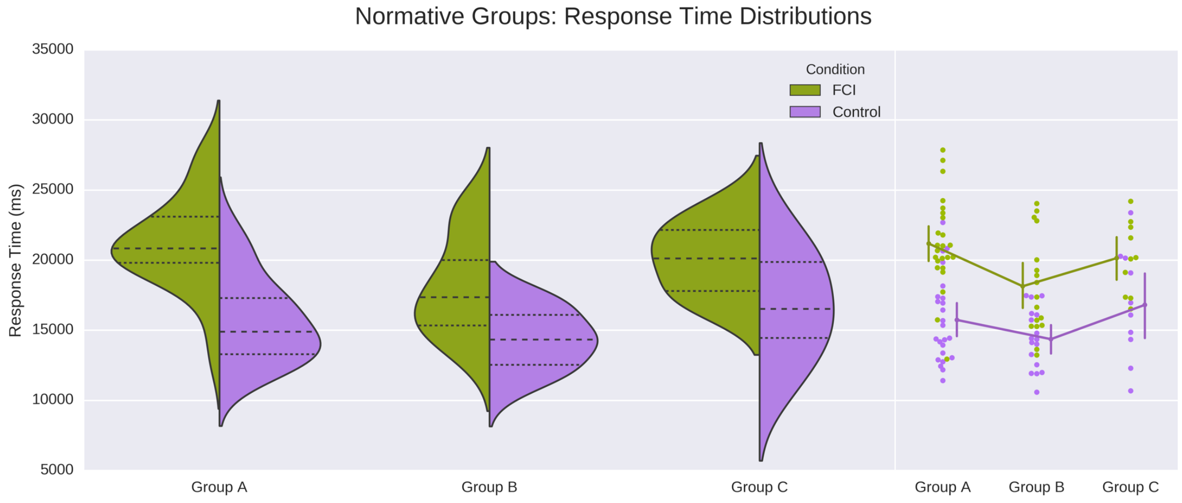


**Fig. S7.** Mean response times (RT; ms) for normative sub-groups across all Phases for FCI and Control conditions. Kernel density estimates of RT distributions are plotted on the left, and the interaction across conditions and groups is provided on the right, with 95% confidence intervals shown as vertical error bars. Significant differences were observed between FCI and Control RTs across all sub-groups ($p << 0.001$). Turkey HSD *post hoc* tests indicated no significant pairwise RT differences in either condition between sub-groups.

RT differences between conditions, phases, and groups were influenced by students' ability to choose when they felt ready to progress to the next phase and when they had finished answering each question. This self-paced task structure emulated that of real-world problem-solving processes and ensured measured brain activity associated with each phase corresponded to intervals in which students were initiating problem solving (Phase I), reading and comprehending the question (Phase II), and choosing an answer (Phase III). While it is possible that RT differences may have impacted the brain activation results we were able to detect, we nonetheless believe allowing for students' authentic and variable problem solving approach was of critical importance in measuring students' problem solving processes. We hold that these RT differences are a central to part of the problem-solving processes.

**Summary of Results.** In this study, we investigated the neural mechanisms underlying physics reasoning across 107 students and identified a fronto-temporo-parietal brain network linked with problem solving. Initiation, question presentation, and answer selection phases evoked integrated V5/MT+, CEN, DAN, and DMN systems. Notably, during answer selection wherein students deliberated between possible outcomes linked to conflicting physics conceptions, they engaged concurrent V5/MT+, lateral fronto-parietal, and DMN activity, evidencing V5 -CEN-DMN engagement as critical for physics reasoning. Follow-up PPI analyses investigating task-based FC between V5/MT+, dlPFC, and RSC found evidence that physics reasoning initiates dorsal stream activity and CEN-DMN information exchange. Strategy and accuracy did not modulate brain activity during reasoning; however, increased difficulty elicited enhanced DAN engagement, representative of reliance on executive functions during demanding problems. Importantly, whole brain activity was not modulated by problem-solving accuracy, but module analysis resulting in a dissociation of student reasoning sub-groups (i.e., problem solvers grouped by similar conceptualizations of physics ideas) yielded ranked performance differences across groups that were linked to conceptual approach. Compellingly, students from groups who applied more Newtonian and coherent physics conceptions showed enhanced engagement of a fronto-temporal network, whereas students who relied on less coherent, non-Newtonian conceptions engaged enhanced visual and SN area activity during problem solving. These insights aid in characterizing the underlying neural processes of how students tackle conflicting physics conceptions during reasoning.

### SUPPLEMENTAL TABLES

**Table S1.** Center of mass activation coordinates for the FCI > Control contrast as reported in MNI space. Cluster region labels are based off those reported by the IBASPM116 Human Brain Atlas.

| Cluster | Hemisphere | Center of Mass (MNI space) | | | Cluster Extent ($mm^3$) | Mean Z Score | Region Labels |
|---|---|---|---|---|---|---|---|
| | | X | Y | Z | | | |
| 1 | B | -34 | 30 | 26 | 85072 | 5.676 | Frontal_Mid_L, Frontal_Sup_L, Frontal_Inf_Tri_L, Frontal_Sup_Medial_L, Precentral_L, Frontal_Inf_Oper_L, Frontal_Inf_Orb_L, Frontal_Mid_Orb_L, Supp_Motor_Area_L, Frontal_Sup_Orb_L, Frontal_Mid_Orb_L, Rolandic_Oper_L, Cingulum_Mid_L, Cingulum_Ant_L, Frontal_Sup_Medial_R, Supp_Motor_Area_R, Temporal_Pole_Sup_L |
| 2 | R | 50 | -50 | 26 | 57088 | 5.222 | Temporal_Inf_R, SupraMarginal_R, Parietal_Inf_R, Temporal_Mid_R, Occipital_Mid_R, Parietal_Sup_R, Angular_R, Postcentral_R, Occipital_Inf_R, Fusiform_R, Precuneus_R, Cerebellum_Crus1_R, Occipital_Sup_R, Cerebellum_6_R |
| 3 | L | -46 | -54 | 42 | 49632 | 6.386 | Parietal_Inf_L, Angular_L, SupraMarginal_L, Parietal_Sup_L, Occipital_Mid_L, Precuneus_L, Postcentral_L, Occipital_Sup_L, Temporal_Sup_L, Temporal_Mid_L |
| 4 | R | 46 | 28 | 18 | 36576 | 4.746 | Frontal_Mid_R, Frontal_Inf_Tri_R, Frontal_Inf_Oper_R, Frontal_Mid_Orb_R, Frontal_Inf_Orb_R, Frontal_Sup_R, Precentral_R, Rolandic_Oper_R, Insula_R |
| 5 | L | -54 | -58 | -8 | 17864 | 5.243 | Temporal_Inf_L, Temporal_Mid_L, Occipital_Inf_L, Occipital_Mid_L, Cerebellum_Crus1_L |
| 6 | R | 28 | -70 | -44 | 13744 | 5.257 | No label generated |
| 7 | L | -32 | -74 | -52 | 6120 | 4.148 | No label generated |
| 8 | L | -8 | -56 | 16 | 1680 | 3.666 | Precuneus_L, Calcarine_L, Cingulum_Post_L, Cuneus_L |
| 9 | L | -12 | 10 | 8 | 1392 | 3.997 | Caudate_L |

**Table S2.** Center of mass activation coordinates for the contrasts (a) FCI Phase I > Control Phase I, (b) FCI Phase II > Control Phase II, and (c) FCI Phase III > Control Phase III as reported in MNI space. Cluster region labels are based off those reported by the IBASPM116 Human Brain Atlas.

**a) FCI Phase I > Control Phase I**

| Cluster | Hemisphere | Center of Mass (MNI space) | | | Cluster Extent ($mm^3$) | Mean Z Score | Labels |
|---|---|---|---|---|---|---|---|
| | | X | Y | Z | | | |
| 1 | B | -16 | -46 | 24 | 295832 | 4.570 | Precentral_L, Parietal_Inf_L, Occipital_Mid_L, Postcentral_L, Frontal_Inf_Tri_L, Frontal_Mid_L, Lingual_R, Calcarine_R, Parietal_Sup_L, Occipital_Mid_R, Calcarine_L, Temporal_Inf_L, Precuneus_L, Parietal_Sup_R, Frontal_Inf_Oper_L, Occipital_Sup_L, Temporal_Mid_L, Parietal_Inf_R, Supp_Motor_Area_L, Lingual_L, Frontal_Sup_L, Cerebelum_Crus1_L, Cerebelum_Crus1_R, Angular_L, SupraMarginal_L, Precuneus_R, Occipital_Sup_R, Fusiform_L, Cerebelum_6_R, Angular_R, Frontal_Sup_Medial_L, Cuneus_R, Occipital_Inf_L, SupraMarginal_R, Cuneus_L, Fusiform_R, Cerebelum_6_L, Cerebelum_Crus2_R, Frontal_Inf_Orb_L, Cerebelum_Crus2_L, Frontal_Mid_Orb_L, Insula_L, Cerebelum_8_R, Postcentral_R, Supp_Motor_Area_R, Rolandic_Oper_L, Cerebelum_7b_R, Cingulum_Mid_L, Cerebelum_7b_L, Frontal_Sup_Medial_R, Cerebelum_9_R, Cingulum_Mid_R, Cerebelum_4_5_R, Cerebelum_8_L, Occipital_Inf_R, Vermis_7, Cingulum_Ant_L, Vermis_8, Frontal_Sup_Orb_L, Vermis_6, Temporal_Mid_R, Temporal_Sup_L, Vermis_4_5 |
| 2 | R | 42 | 12 | 36 | 26408 | 3.970 | Frontal_Mid_R, Frontal_Inf_Oper_R, Frontal_Inf_Tri_R, Precentral_R, Frontal_Sup_R, Rolandic_Oper_R |
| 3 | R | 56 | -54 | -16 | 11680 | 4.148 | Temporal_Inf_R, Temporal_Mid_R, Cerebelum_Crus1_R, Fusiform_R, Occipital_Inf_R |
| 4 | B | -10 | -24 | -4 | 11048 | 3.788 | Thalamus_L, Hippocampus_L, ParaHippocampal_L, Lingual_L, Thalamus_R, Precuneus_L, |

| | | | | | | | |
|---|---|---|---|---|---|---|---|
| | | | | | | | Cerebelum_4_5_L, Vermis_3, Pallidum_L, Amygdala_L |
| 5 | B | -4 | -26 | 26 | 5664 | 3.862 | Cingulum_Post_L, Cingulum_Mid_L, Precuneus_L, Cingulum_Ant_L, Cingulum_Mid_R, Calcarine_L |
| 6 | R | 34 | 24 | -8 | 2280 | 4.186 | Insula_R, Frontal_Inf_Orb_R |
| 7 | L | -32 | -72 | -60 | 1408 | 3.814 | |
| **b) FCI Phase II < Control Phase II** | | | | | | | |
| 1 | R | 40 | -56 | 32 | 43064 | 4.630 | Occipital_Mid_R, Parietal_Sup_R, Temporal_Inf_R, Postcentral_R, Parietal_Inf_R, SupraMarginal_R, Occipital_Inf_R, Precuneus_R, Fusiform_R, Occipital_Sup_R, Temporal_Mid_R, Angular_R, Cerebelum_Crus1_R, Cerebelum_6_R |
| 2 | L | -34 | -64 | 32 | 30072 | 4.405 | Occipital_Mid_L, Parietal_Sup_L, Parietal_Inf_L, Occipital_Inf_L, Precuneus_L, Temporal_Inf_L, SupraMarginal_L, Occipital_Sup_L, Temporal_Mid_L, Postcentral_L, Angular_L |
| 3 | L | -26 | -2 | 54 | 8344 | 5.003 | Frontal_Mid_L, Frontal_Sup_L, Precentral_L |
| 4 | L | -48 | 40 | 4 | 6760 | 3.888 | Frontal_Inf_Tri_L, Frontal_Inf_Orb_L, Frontal_Mid_Orb_L, Frontal_Mid_L |
| 5 | R | 26 | -2 | 52 | 6528 | 4.770 | Frontal_Sup_R, Frontal_Mid_R, Precentral_R |
| 6 | R | 52 | 10 | 20 | 3168 | 4.3639 | Frontal_Inf_Oper_R, Precentral_R, Rolandic_Oper_R, Frontal_Inf_Tri_R |
| 7 | L | -50 | 6 | 20 | 1464 | 3.670 | Frontal_Inf_Oper_L, Precentral_L, Rolandic_Oper_L |
| **c) FCI Phase III < Control Phase III** | | | | | | | |
| 1 | L | -54 | -56 | 26 | 50152 | 5.606 | Temporal_Mid_L, Angular_L, Parietal_Inf_L, SupraMarginal_L, Occipital_Mid_L, Temporal_Inf_L, Temporal_Sup_L, Parietal_Sup_L, Occipital_Inf_L |
| 2 | B | -14 | 42 | 32 | 50128 | 4.809 | Frontal_Sup_L, Frontal_Sup_Medial_L, Frontal_Mid_L, Frontal_Mid_Orb_L, Frontal_Sup_Medial_R, Frontal_Mid_Orb_R, Supp_Motor_Area_L, Frontal_Sup_R, Frontal_Sup_Orb_L, Precentral_L, Rectus_L, Frontal_Sup_Orb_R, Rectus_R, Frontal_Mid_Orb_L, Supp_Motor_Area_R, Frontal_Inf_Oper_L |
| 3 | R | 58 | -54 | 26 | 27744 | 4.478 | Angular_R, Temporal_Mid_R, SupraMarginal_R, Parietal_Inf_R, Temporal_Inf_R, Occipital_Mid_R, Temporal_Sup_R, Postcentral_R |

| | | | | | | | |
|---|---|---|---|---|---|---|---|
| 4 | L | -48 | 36 | -10 | 21792 | 4.617 | Frontal_Inf_Orb_L, Frontal_Inf_Tri_L, Frontal_Mid_Orb_L, Frontal_Inf_Oper_L, Frontal_Mid_L, Temporal_Pole_Sup_L |
| 5 | B | -4 | -48 | 26 | 20344 | 4.496 | Precuneus_L, Cingulum_Post_L, Cingulum_Mid_L, Cuneus_L, Cingulum_Post_R, Precuneus_R, Cingulum_Mid_R, Calcarine_L, Cerebelum_4_5_L, Lingual_L, Vermis_4_5, Cingulum_Ant_L |
| 6 | R | 28 | -78 | -44 | 11680 | 4.961 | |
| 7 | R | 54 | 34 | -4 | 10160 | 3.929 | Frontal_Inf_Tri_R, Frontal_Inf_Orb_R, Frontal_Inf_Oper_R, Frontal_Mid_Orb_R, Frontal_Mid_R |
| 8 | L | -30 | -80 | -50 | 5560 | 3.823 | |
| 9 | L | -12 | 10 | 12 | 1240 | 3.877 | Caudate_L |
| 10 | L | -14 | -8 | 18 | 64 | 3.214 | Caudate_L, Thalamus_L |

**Table S3.** Meta-analytic functional decoding for unthresholded z-statistic (a) FCI > Control (all phases), (b) FCI Phase I > Control Phase I, (c) FCI Phase II > Control Phase II, (d) FCI Phase III > Control Phase III, and (e) problem difficulty modulator maps. Decoding was performed with a 200-topic GC-LDA (4) topic model trained on the Neurosynth database. The top ten terms returned are provided along with their associated Neurosynth correlation values.

| **a) Full Questions** | |
|---|---|
| **Term** | **Weight** |
| switching | 309.71084 |
| default | 276.1635173 |
| motion | 252.5753468 |
| reasoning | 214.5672386 |
| gestures | 211.2976516 |
| ambiguous | 180.1752178 |
| default_mode | 147.8986879 |
| switch | 137.3589833 |
| body | 135.9365735 |
| relational | 117.1724557 |
| **b) Phase I: Problem Initiation** | |
| **Term** | **Weight** |
| visual | 6280.671847 |
| motor | 3552.329718 |
| spatial | 2180.073879 |
| attention | 1794.573378 |
| memory | 1503.498075 |
| perceptual | 1334.731314 |
| working_memory | 1212.675255 |
| words | 1171.632954 |
| retrieval | 1047.038397 |
| sensory | 1044.932068 |
| **c) Phase II: Question Presentation** | |
| **Term** | **Weight** |
| switching | 174.7799387 |
| visuo | 143.0582605 |
| grasping | 95.3737437 |
| switch | 83.24127548 |
| vstm | 80.24913636 |
| numbers | 80.15246068 |
| numerical | 78.16072276 |
| drawing | 68.03863533 |
| grasp | 63.52682393 |
| hands | 62.93979874 |
| **d) Phase III: Answer Selection** | |
| **Term** | **Weight** |
| default | 649.6765185 |
| default_mode_network | 447.7949888 |
| mentalizing | 329.1184083 |
| default_mode | 240.3050669 |
| intrinsic | 215.4858992 |
| mental_states | 186.8147481 |
| seed | 150.9431503 |

| | |
|---|---|
| free | 124.7647576 |
| mind | 121.8747808 |
| ambiguous | 111.3774804 |
| **e) Difficulty Modulator** | |
| **Term** | **Weight** |
| visual | 4998.611373 |
| spatial | 2165.71031 |
| motion | 1454.923421 |
| motor | 1451.786037 |
| face | 1361.697261 |
| perceptual | 1270.365251 |
| body | 1211.229165 |
| faces | 1162.215922 |
| perception | 1046.94595 |
| memory | 785.0733801 |

**Table S4.** Center of mass coordinates for psychophysiological interaction (PPI) task-based functional connectivity between dlPFC, RSC, and V5/MT+ seeds associated with the contrast FCI > Control, as reported in MNI space. Cluster region labels are based off those reported by the IBASPM116 Human Brain Atlas.

| Cluster | Hemisphere | Center of Mass (MNI space) | | | Cluster Extent ($mm^3$) | Mean Z Score | Labels |
|---|---|---|---|---|---|---|---|
| | | X | Y | Z | | | |
| **a) Left V5/MT+ Seed** | | | | | | | |
| 1 | R | 54 | -30 | 40 | 14128 | 4.104 | SupraMarginal_R, Postcentral_R, Parietal_Inf_R, Parietal_Sup_R, Rolandic_Oper_R |
| 2 | R | 50 | -62 | -8 | 11176 | 4.334 | Temporal_Inf_R, Temporal_Mid_R, Occipital_Inf_R, Fusiform_R, Occipital_Mid_R |
| 3 | L | -48 | -72 | -2 | 5344 | 4.131 | Occipital_Mid_L, Occipital_Inf_L, Temporal_Mid_L, Temporal_Inf_L |
| 4 | R | 24 | -70 | 46 | 2216 | 3.607 | Occipital_Sup_R, Parietal_Sup_R, Precuneus_R, Occipital_Mid_R, Angular_R, Cuneus_R |
| 5 | L | -60 | -24 | 34 | 1776 | 3.386 | SupraMarginal_L, Parietal_Inf_L, Postcentral_L |
| 6 | L | -32 | -54 | 54 | 1256 | 3.369 | Parietal_Inf_L, Parietal_Sup_L |
| **b) Left dlPFC Seed** | | | | | | | |
| 1 | R | 46 | -60 | 0 | 39504 | 4.181 | Temporal_Mid_R, Temporal_Inf_R, Occipital_Mid_R, Fusiform_R, Occipital_Inf_R, Cerebelum_Crus1_R, Temporal_Sup_R, Cerebelum_4_5_R, Angular_R, Cerebelum_6_R, SupraMarginal_R, ParaHippocampal_R |
| 2 | L | -44 | -72 | 4 | 28432 | 4.200 | Occipital_Mid_L, Temporal_Mid_L, Fusiform_L, Occipital_Inf_L, Temporal_Inf_L, Cerebelum_6_L, Angular_L, Cerebelum_4_5_L, Parietal_Inf_L |
| 3 | B | 4 | -50 | 24 | 8096 | 3.489 | Precuneus_R, Precuneus_L, Cingulum_Post_L, Cingulum_Mid_R, Cingulum_Post_R, Cingulum_Mid_L, Vermis_4_5, Calcarine_R, Cuneus_L, Cuneus_R, Lingual_R |
| 4 | B | 4 | 62 | -2 | 4168 | 3.517 | Frontal_Sup_Medial_R, Frontal_Mid_Orb_L, Frontal_Mid_Orb_R, Frontal_Sup_Medial_L, Frontal_Sup_R |
| 5 | R | 58 | -24 | 40 | 1672 | 3.631 | SupraMarginal_R, Postcentral_R |
| **c) Left RSC Seed** | | | | | | | |
| 1 | B | 0 | 36 | 22 | 23072 | 3.727 | Frontal_Sup_Medial_L, |

| | | | | | | | |
|---|---|---|---|---|---|---|---|
| | | | | | | | Supp_Motor_Area_L, Cingulum_Ant_R, Cingulum_Ant_L, Frontal_Mid_Orb_R, Frontal_Sup_Medial_R, Cingulum_Mid_R, Frontal_Mid_Orb_L, Cingulum_Mid_L, Supp_Motor_Area_R, Frontal_Sup_L, Rectus_R |
| 2 | L | -32 | -64 | -18 | 12432 | 3.972 | Fusiform_L, Cerebelum_6_L, Cerebelum_Crus1_L, Lingual_L, Temporal_Inf_L, Occipital_Inf_L, Cerebelum_4_5_L, Calcarine_L |
| 3 | L | -28 | -70 | 36 | 7952 | 4.007 | Occipital_Mid_L, Parietal_Sup_L, Parietal_Inf_L, Occipital_Sup_L, Angular_L |
| 4 | L | -44 | 12 | 26 | 4512 | 3.499 | Frontal_Inf_Tri_L, Precentral_L, Frontal_Inf_Oper_L, Postcentral_L, Rolandic_Oper_L |
| 5 | B | 0 | -34 | 28 | 3832 | 3.708 | Cingulum_Mid_L, Cingulum_Post_L, Cingulum_Mid_R, Cingulum_Post_R |
| 6 | L | -56 | -32 | -6 | 3480 | 3.798 | Temporal_Mid_L |
| 7 | R | 30 | -64 | 44 | 2816 | 3.588 | Angular_R, Parietal_Sup_R, Occipital_Sup_R, Occipital_Mid_R, Parietal_Inf_R |
| 8 | R | 30 | 44 | 26 | 1944 | 3.563 | Frontal_Mid_R, Frontal_Sup_R |
| 9 | R | 38 | 16 | -2 | 1296 | 3.620 | Insula_R, Frontal_Inf_Orb_R, Frontal_Inf_Oper_R |
| 10 | B | -2 | -50 | 10 | 1104 | 3.570 | Precuneus_L, Cingulum_Post_L, Vermis_4_5, Cingulum_Post_R, Calcarine_L, Lingual_L |
| 11 | R | 30 | 22 | -20 | 1104 | 3.654 | Frontal_Inf_Orb_R, Insula_R, Temporal_Pole_Sup_R, Temporal_Pole_Mid_R |

**Table S5.** Activation coordinates associated with the whole-brain parametric modulation of FCI > Control (all phases) by normative problem difficulty (2), as reported in MNI space. Cluster region labels are based off those reported by the IBASPM116 Human Brain Atlas.

| **Whole-Brain Activity Correlated with Problem Difficulty** | | | | | | | |
|---|---|---|---|---|---|---|---|
| Cluster | Hemisphere | Center of Mass (MNI space) | | | Cluster Extent ($mm^3$) | Mean Z Score | Labels |
| | | X | Y | Z | | | |
| 1 | B | 38 | -60 | 16 | 118720 | 5.750 | Occipital_Mid_R, Fusiform_R, Parietal_Sup_R, Temporal_Inf_R, Temporal_Mid_R, Cerebelum_6_R, SupraMarginal_R, Precuneus_R, Postcentral_R, Occipital_Inf_R, Occipital_Sup_R, Parietal_Inf_R, Cerebelum_Crus1_R, Angular_R, Cerebelum_4_5_R, Cuneus_R, Lingual_R, Calcarine_R, ParaHippocampal_R, Rolandic_Oper_R, Cerebelum_Crus2_R, Precuneus_L |
| 2 | L | -36 | -62 | 16 | 95296 | 5.539 | Occipital_Mid_L, Parietal_Inf_L, Parietal_Sup_L, Fusiform_L, Occipital_Inf_L, Cerebelum_6_L, Temporal_Inf_L, Cerebelum_Crus1_L, Occipital_Sup_L, SupraMarginal_L, Temporal_Mid_L, Precuneus_L, Postcentral_L, Cerebelum_8_L, Cerebelum_9_L, Cerebelum_4_5_L, Cerebelum_Crus2_L, Angular_L, Cerebelum_7b_L, Lingual_L, Cuneus_L |
| 3 | R | 40 | 4 | 34 | 21960 | 4.593 | Precentral_R, Frontal_Inf_Oper_R, Frontal_Sup_R, Frontal_Mid_R, Insula_R, Rolandic_Oper_R, Frontal_Inf_Tri_R, Putamen_R, Supp_Motor_Area_R, Temporal_Pole_Sup_R |
| 4 | L | -26 | -6 | 54 | 7560 | 4.714 | Precentral_L, Frontal_Sup_L, Frontal_Mid_L, Supp_Motor_Area_L |
| 5 | L | -20 | -72 | -56 | 3104 | 4.079 | |
| 6 | L | -54 | 4 | 28 | 2848 | 4.192 | Precentral_L, Frontal_Inf_Oper_L, Rolandic_Oper_L |
| 7 | R | 22 | -48 | -58 | 2760 | 3.873 | |
| 8 | R | 46 | 38 | 6 | 1752 | 3.721 | Frontal_Inf_Tri_R, Frontal_Mid_R |
| 9 | L | -40 | -2 | 6 | 1352 | 4.289 | Insula_L, Rolandic_Oper_L |
| 10 | R | 18 | -30 | -6 | 848 | 3.440 | Thalamus_R, ParaHippocampal_R, Hippocampus_R, Lingual_R |

**Table S6.** Conceptual modules, their constituent FCI answer choices, and their descriptions, as taken from (14). Bolded FCI answer choices represent the items that we adapted for in-scanner presentation. In cases where (14) did not describe student conceptualizations associated with a module, we have provided additional interpretations that add/expand upon the original descriptions to aid in interpretation of outcomes in the present study. Modules that we have elaborated upon are marked with an obelisk †. Additionally, where appropriate we provide external references that describe further observations of the common incorrect physical conceptions detailed by a conceptual module.

**Common Non-Newtonian Conceptual Modules** (14)

| | **Module** | **Constituent FCI Answer Choices** | **Detailed Conceptual Description** |
|---|---|---|---|
| m1 | Moving objects experience an “impetus” force | **2B**, **3B**, 5E, **6A**, **7A**, **7E**, **8D**, 8E, 13C, **14A**, **14C**, 17D, 18E, 19B, 20A, 21A, 22D, 23D, 23E, 24C, 24D, 25A, 25B, 25E, 26A, **27B**, 30E | If an object is moving, then there must be a force actively causing the motion. This fictional force is referred to as an “impetus” force. Students who hold this view often similarly believe that, if an object’s motion becomes diminished, a diminishing impetus force must have caused the change. This (incorrect) Galilean model was held by many medieval physicists and is often characterized by a common confusion among students between the concepts of force and velocity. The impetus force fallacy is a prevalent, particularly coherent, and persistent model that students’ apply across contextually diverse situations (15, 16). |
| m2 | More force yields more result | 1A, **2C**, 2E, **3E**, 4A, 10C, 11E, 15C, 19C, 20E, 21D, 26B, 26C, 26D, **27D**, 28D, 30A | If a force acts on an object and is increased, then *something* about the object’s motion must also increase. This idea is correct if the increased quantity is acceleration. However, students holding this view often assign the increased quantity incorrectly and/or without justification (e.g., displacement, time, velocity, or an additional non-physical force). Similarly, how the quantity increases (e.g., constantly or scaled by a factor) is often assigned incorrectly and/or without justification. Because of the vague/unstructured nature of what quantity increases and how, this module is applied in different and sometimes conflicting ways depending on the problem or context. This module is compatible with the physics phenomenological primitive, or *p-prim*, known as “Ohm’s p-prim” (19). P-prims are irreducible, loosely connected sets of intuitive physics knowledge that students use to explain physical phenomena (19). Ohm’s p-prim asserts that how much result something receives is proportional to the amount of resistance it gives (more effort implies more result and more resistance implies less result). In contrast to more stable and consistent sets of physics ideas, such as those describe in *m1*, p-prims such as the one paralleled in this module illustrate more fragmentary intuitive knowledge pieces that describes contextually situated emergent knowledge that student’s use when reasoning (16). |
| m3 | Competing forces cause motion, or | 4D, **6C**, 11B, 16C, 17A, 20B, 20C, 25D, 28C | This module is described by two competing interpretations: 1) competing forces cause motion (e.g., motion occurs because one force “wins” out over another competing |

| | | | |
|---|---|---|---|
| | acceleration and velocity are not distinguished | | force), and/or 2) students fail to discriminate between velocity and acceleration, thus a net force yields a velocity. |
| m4 | A moving object's impetus eventually "runs out"† | 5D, **8E**, 10D, 11C, 15D, 16D, 18D | This module is likely a variant of the impetus force module (*m1*). However, the ways in which this module varies from *m1* is not specified in the original paper (14). We interpret this module as representing an impetus conception of force wherein a moving objects' impetus force "runs out" over time. This is characterized by the belief that objects have a natural tendency to remain still. That is, students who hold this view may believe objects set in motion by an active agent stores the external force as it moves, but then releases its impetus over time due to a natural tendency of all objects to remain inactive. |
| m5 | Confusion in relating an object's speed and path† | 5C, 9C, **12C**, **12D**, 13B, 18C, 19A, 22C, **27A** | This module is less consistent, and therefore less characterized by concrete, coherent non-physical beliefs. In (14), the module is described as indicating an indistinct lack of understanding about velocity. We further characterize it here as students reaching an incorrect conclusion that involves relating a moving object's path to its speed. However, the way in which students relate path to speed is not applied consistently, indicating students do not have a clear strategy and may be confused. |
| m6 | A sudden force on an object induces an instantaneous path change | **7C**, **8A**, 9B, 15E, 16E, 17E, 21B, 23C, 28A | The module is characterized by the belief that when a moving object undergoes a quick change in force it will instantaneously (e.g., over an infinitesimally small time interval) alter its path to move in the direction of the external force. |
| m7 | An object's mass determines how it falls | 1D, **2D**, 9D, 10E, 18A, 19D, 23A | The incorrect belief that objects of different masses fall at different rates and traverse different horizontal distances as they fall. This view is frequently compatible with the Aristotelian view of falling bodies wherein more massive objects fall faster, although there is evidence that student's naïve conceptions about how mass relates to trajectory don't share the same coherence as Aristotle's description (23). FCI answer choices in this module suggest students may view this supposed mass to time of flight/distance relationship as not linearly related. This module may be an iteration of the *more force yields more result* module (*m2*). |
| m8 | Indistinct confusion regarding downward force or scenario description† | **14B**, 21C, 22A, **29D** | This model was originally proposed as reflecting student confusion with interpreting the physical scenario described in a particular FCI question (14). However, because the module is composed of answer choices not related to a single FCI question, we have extended this interpretation to describe an indistinct confusion about either the scenario descriptions or downward force. Two answer choices in this module (21C, 22A) indicate students may be confused about |

| | | | |
|---|---|---|---|
| | | | the physical scenario described in one FCI question (in particular the length of time a force is applied to an object). An additional item (29D) indicates students believe air exerts a dominant downward force on objects, while another item (14B) indicates students believe objects fall vertically even after being released with an initial horizontal velocity. Thus, this module involves disjoint ideas that are difficult to characterize as a single coherent conceptual structure. We interpret it as involving unidentifiable confusions about force and/or the physical description of a question. |
| m9 | Confusion regarding gravitational action† | 1B, **3A**, 5A, 11A, 28B, 29C, 30B | Multiple answer items in this set (1B, 3A, 5A, 11A, 30B) share a focus on the gravitational force as acting in replacement of, or dominant to other forces. Thus, the module is described in terms of gravity as being a constant factor in each incorrect answer (14). We additionally observe that this confusion about gravitational action appears to impact how students predict resulting motion or itemize which forces act on moving and/or stationary objects. How gravity impacts motion and/or free body diagrams differs from answer to answer, indicating this module may represent somewhat inconsistent ideas about gravity. |

**Table S7.** Center of mass coordinates associated, as reported in MNI space, of brain activation (FCI > Control, all Phases) group differences between normative sub-groups identified by module analysis of student answer distributions. Omnibus test results are listed in (a) and (b), with F-scores converted to z statistics. Results from *post hoc* t-tests investigating differences across each pair of sub-groups are listed in (c)-(h). Cluster region labels are based off those reported by the IBASPM116 Human Brain Atlas.

| Cluster | Hemisphere | Center of Mass (MNI space) | | | Cluster Extent ($mm^3$) | Mean Z Score | Labels |
|---|---|---|---|---|---|---|---|
| | | X | Y | Z | | | |
| **a) Whole-brain one-way ANOVA: Group A or B vs. Group C** | | | | | | | |
| 1 | B | 0 | -78 | 6 | 12384 | 3.627 | Calcarine_L, Lingual_L, Calcarine_R, Cuneus_R, Cuneus_L, Lingual_R, Occipital_Sup_L, Cerebelum_6_L, Precuneus_R |
| 2 | L | -48 | -18 | 54 | 4728 | 3.504 | Postcentral_L, Precentral_L, Parietal_Inf_L |
| 3 | L | -8 | 10 | 36 | 1608 | 3.621 | Cingulum_Mid_L, Cingulum_Ant_L, Supp_Motor_Area_L |
| 4 | L | -44 | 50 | -4 | 1592 | 3.534 | Frontal_Mid_Orb_L, Frontal_Mid_L, Frontal_Inf_Orb_L, Frontal_Inf_Tri_L, Frontal_Sup_L |
| **b) Whole-brain one-way ANOVA: Group A or C vs. Group B** | | | | | | | |
| 1 | B | 0 | -78 | 6 | 12368 | 3.627 | Calcarine_L, Lingual_L, Calcarine_R, Cuneus_R, Cuneus_L, Lingual_R, Occipital_Sup_L, Cerebelum_6_L, Precuneus_R |
| 2 | L | -48 | -18 | 54 | 4720 | 3.504 | Postcentral_L, Precentral_L, Parietal_Inf_L |
| 3 | L | -8 | 10 | 36 | 1608 | 3.620 | Cingulum_Mid_L, Cingulum_Ant_L, Supp_Motor_Area_L |
| 4 | L | -44 | 50 | -4 | 1592 | 3.533 | Frontal_Mid_Orb_L, Frontal_Mid_L, Frontal_Inf_Orb_L, Frontal_Inf_Tri_L, Frontal_Sup_L |
| **c) Group A > Group B** | | | | | | | |
| No significant group differences detected | | | | | | | |
| **d) Group B > Group A** | | | | | | | |
| No significant group differences detected | | | | | | | |
| **e) Group A > Group C** | | | | | | | |
| 2 | L | -44 | 48 | -2 | 2592 | 3.409 | Frontal_Mid_Orb_L, Frontal_Inf_Tri_L, Frontal_Mid_L, Frontal_Inf_Orb_L |
| 3 | R | 38 | -70 | -50 | 2400 | 3.356 | |
| 4 | L | -58 | -62 | -4 | 1752 | 3.418 | Temporal_Mid_L, Temporal_Inf_L, Occipital_Inf_L, Occipital_Mid_L |
| 5 | R | 60 | -50 | -12 | 1464 | 3.409 | Temporal_Inf_R, Temporal_Mid_R |
| **f) Group C > Group A** | | | | | | | |
| 1 | B | -2 | -80 | 6 | 14232 | 3.673 | Calcarine_L, Lingual_L, Calcarine_R, Cuneus_R, Cuneus_L, Lingual_R, |

| | | | | | | | |
|---|---|---|---|---|---|---|---|
| | | | | | | | Occipital_Inf_L, Occipital_Mid_L, Fusiform_L, Cerebelum_6_L, Cerebelum_Crus1_L, Occipital_Sup_L |
| 2 | L | -46 | -18 | 54 | 9496 | 3.643 | Postcentral_L, Precentral_L, Parietal_Inf_ |
| 3 | B | 0 | 10 | 42 | 6640 | 3.522 | Supp_Motor_Area_R, Cingulum_Mid_L, Cingulum_Mid_R, Supp_Motor_Area_L, Cingulum_Ant_L, Cingulum_Ant_R, Frontal_Sup_R |
| 4 | R | 46 | -12 | 52 | 3648 | 3.464 | Precentral_R, Frontal_Mid_R, Postcentral_R |
| 5 | R | 46 | 12 | -10 | 2688 | 3.640 | Insula_R, Temporal_Pole_Sup_R, Frontal_Inf_Orb_R |
| 6 | L | -48 | 8 | -8 | 1840 | 3.410 | Temporal_Pole_Sup_L, Insula_L, Temporal_Mid_L, Temporal_Sup_L, Rolandic_Oper_L |
| 7 | L | -56 | -24 | 8 | 1680 | 3.414 | Temporal_Sup_L, Temporal_Mid_L, Postcentral_L, Heschl_L, Rolandic_Oper_L |
| **g) Group B > Group C** | | | | | | | |
| 1 | L | -48 | 48 | -6 | 1328 | 3.465 | Frontal_Mid_Orb_L, Frontal_Inf_Orb_L, Frontal_Inf_Tri_L, Frontal_Mid_L |
| **h) Group C > Group B** | | | | | | | |
| 1 | B | 2 | -72 | 8 | 10664 | 3.448 | Lingual_L, Lingual_R, Calcarine_L, Cuneus_R, Calcarine_R, Precuneus_R, Vermis_6, Cuneus_L, Cerebelum_4_5_L, Vermis_4_5, Cerebelum_6_L, Cerebelum_6_R |
| 2 | L | -20 | -50 | -8 | 256 | 3.243 | Lingual_L, Fusiform_L |
| 3 | R | 26 | -74 | 6 | 40 | 3.175 | No label generated |

**Table S8.** A significant interaction was detected in the two-way mixed effects ANOVA of Condition of (FCI, Control) x Phase (Phase I, Phase II, Phase III) on RT across all 107 students. Results from follow up Turkey HSD *post hoc* tests on the family of six estimates are provided below.

| Contrast | Estimate | SE | df | t ratio | P value |
|---|---|---|---|---|---|
| Phase I Control - Phase II Control | 3966.642 | 198.0183 | 535 | 20.032 | <.0001 |
| Phase I Control - Phase III Control | 592.2531 | 198.0183 | 535 | 2.991 | 0.0344 |
| Phase I Control - Phase I FCI | 329.6553 | 198.0183 | 535 | 1.665 | 0.5558 |
| Phase I Control - Phase II FCI | 1633.9599 | 198.0183 | 535 | 8.252 | <.0001 |
| Phase I Control - Phase III FCI | -1850.9198 | 198.0183 | 535 | -9.347 | <.0001 |
| Phase II Control - Phase III Control | -3374.3889 | 198.0183 | 535 | -17.041 | <.0001 |
| Phase II Control - Phase I FCI | -3636.9866 | 198.0183 | 535 | -18.367 | <.0001 |
| Phase II Control - Phase II FCI | -2332.6821 | 198.0183 | 535 | -11.78 | <.0001 |
| Phase II Control - Phase III FCI | -5817.5617 | 198.0183 | 535 | -29.379 | <.0001 |
| Phase III Control - Phase I FCI | -262.5977 | 198.0183 | 535 | -1.326 | 0.7704 |
| Phase III Control - Phase II FCI | 1041.7068 | 198.0183 | 535 | 5.261 | <.0001 |
| Phase III Control - Phase III FCI | -2443.1728 | 198.0183 | 535 | -12.338 | <.0001 |
| Phase I FCI - Phase II FCI | 1304.3045 | 198.0183 | 535 | 6.587 | <.0001 |
| Phase I FCI - Phase III FCI | -2180.5751 | 198.0183 | 535 | -11.012 | <.0001 |
| Phase II FCI - Phase III FCI | -3484.8796 | 198.0183 | 535 | -17.599 | <.0001 |
